# Human iPSC-derived RPE and retinal organoids reveal impaired alternative splicing of genes involved in pre-mRNA splicing in PRPF31 autosomal dominant retinitis pigmentosa

**DOI:** 10.1101/232397

**Authors:** Adriana Buskin, Lili Zhu, Valeria Chichagova, Basudha Basu, Sina Mozaffari-Jovin, David Dolan, Alastair Droop, Joseph Collin, Revital Bronstein, Sudeep Mehrotra, Michael Farkas, Gerrit Hilgen, Kathryn White, Dean Hallam, Katarzyna Bialas, Git Chung, Carla Mellough, Yuchun Ding, Natalio Krasnogor, Stefan Przyborski, Jumana Al-Aama, Sameer Alharthi, Yaobo Xu, Gabrielle Wheway, Katarzyna Szymanska, Martin McKibbin, Chris F Inglehearn, David J Elliott, Susan Lindsay, Robin R Ali, David H Steel, Lyle Armstrong, Evelyne Sernagor, Eric Pierce, Reinhard Lüehrmann, Sushma-Nagaraja Grellscheid, Colin A Johnson, Majlinda Lako

## Abstract

Mutations in pre-mRNA processing factors (PRPFs) cause 40% of autosomal dominant retinitis pigmentosa (RP), but it is unclear why mutations in ubiquitously expressed PRPFs cause retinal disease. To understand the molecular basis of this phenotype, we have generated RP type 11 *(PRPF31-*mutated) patient-specific retinal organoids and retinal pigment epithelium (RPE) from induced pluripotent stem cells (iPSC). Impaired alternative splicing of genes encoding pre-mRNA splicing proteins occurred in patient-specific retinal cells and *Prpf31^+/−^* mouse retinae, but not fibroblasts and iPSCs, providing mechanistic insights into retinal-specific phenotypes of PRPFs. RPE was the most affected, characterised by loss of apical-basal polarity, reduced trans-epithelial resistance, phagocytic capacity, microvilli, and cilia length and incidence. Disrupted cilia morphology was observed in patient-derived-photoreceptors that displayed progressive features associated with degeneration and cell stress. *In situ* gene-editing of a pathogenic mutation rescued key structural and functional phenotypes in RPE and photoreceptors, providing proof-of-concept for future therapeutic strategies.

**eTOC:** PRPF31 is a ubiquitously expressed pre-mRNA processing factor that when mutated causes autosomal dominant RP. Using a patient-specific iPSC approach, Buskin and Zhu *et al.* show that retinal-specific defects result from altered splicing of genes involved in the splicing process itself, leading to impaired splicing, loss of RPE polarity and diminished phagocytic ability as well as reduced cilia incidence and length in both photoreceptors and RPE.

**Highlights:**

- Successful generation of iPSC-derived RPE and photoreceptors from four RP type 11 patients
- RPE cells express the mutant PRPF31 protein and show the lowest expression of wildtype protein
- *PRPF31* mutations result in altered splicing of genes involved in pre-mRNA splicing in RPE and retinal organoids
- *Prpf31* haploinsufficiency results in altered splicing of genes involved in pre-mRNA splicing in mouse retina
- RPE cells display loss of polarity, reduced barrier function and phagocytosis
- Photoreceptors display shorter and fewer cilia and degenerative features
- RPE cells display most abnormalities suggesting they might be the primary site of pathogenesis
- *In situ* gene editing corrects the mutation and rescues key phenotypes

## Introduction

Retinitis pigmentosa (RP) is one of the most common inherited forms of retinal blindness with a prevalence of about 1 in 2500 births and more than 1 million people affected worldwide (Hartong et al., 2006). RP is characterized by progressive degeneration of the mid-peripheral retina, leading to night blindness, visual field constriction, and eventual loss of visual acuity. To date, there are no effective treatments for RP and it remains a medically challenging disease.

Autosomal dominant inheritance accounts for about 40% of RP, of which an estimated 38% are caused by mutations in seven pre-mRNA processing factors (PRPFs) *PRPF8, PRPF31, PRPF3, PRPF4, PRPF6, SNRNP200* and *RP9* (Chakarova et al., 2002; Chen et al., 2014; Ezquerra-Inchausti et al., 2017; Keen et al., 2002; Maita et al., 2000; McKie et al., 2001; Sullivan et al., 2006; Tanackovic et al., 2011a; Vithana et al., 2001; Zhao et al., 2009). All seven PRPFs are components of the U4/U6.U5 tri-snRNP (small nuclear ribonucleoprotein) subunit of the spliceosome, the large RNP complex that catalyses pre-mRNA splicing. Paradoxically, PRPFs are ubiquitously expressed (Tanackovic et al., 2011a), but mutations only cause retinal-specific degeneration, raising the question of why retinal cells are more susceptible to deficiencies in these splicing factors.

Previous work has indicated that mutations in PRPFs affect the stoichiometry and kinetics of spliceosome assembly (Tanackovic et al., 2011b; Tanackovic and Rivolta, 2009), resulting in either transcriptional dysregulation of key genes needed for retinal function (Ivings et al., 2008), or mis-folding and aggregation of mutant PRPF proteins in photoreceptors that triggers apoptosis (Comitato et al., 2007; Ivings et al., 2008). Recently, an unbiased reverse-genetics screen carried out by us has also shown that several PRPFs (PRPF31, PRPF8 and PRPF6) localize specifically to the basal body complex and connecting cilium of photoreceptors (Wheway et al., 2015). Despite this progress, the disease mechanisms for PRPF-related RPs remain unclear and it is uncertain whether retinal pigment epithelium (RPE) or photoreceptors are the primary affected tissue. PRPF animal models are available, but some do not have an RP phenotype (Bujakowska et al., 2009; Graziotto et al., 2008), whilst others only display late-onset defects in the RPE (Farkas et al., 2014; Graziotto et al., 2011).

Alternative pre-mRNA splicing expands the coding capacity of eukaryotic genomes by differential inclusion of exons or retention of introns in mRNA that enables a relatively small number of genes to encode a diverse proteome. High levels of splicing diversity occur in the vertebrate nervous system where it is required for neuronal development and function. Mouse rod and cone photoreceptors have a specific splicing programme that is initiated prior to the development of outer segments and appears to be driven by the Musashi protein (Murphy et al., 2016). This specific splicing programme primarily affects transcripts encoding components of photoreceptor primary cilia and outer segments, both of which are essential for phototransduction. Strikingly, the splicing programme utilizes "retinal-specific exons" that are included for photoreceptor transcripts but are excluded in non-retinal cell-types. Collectively, these data suggest that precise regulation of splicing is critically important for generation of photoreceptor-specific transcripts including those that are involved in ciliogenesis (Murphy et al., 2016).

Induced pluripotent stem cells (iPSC) (Takahashi et al., 2007), can be guided to differentiate into any tissue type including photoreceptor and RPE cells (Carr et al., 2009; Kuwahara et al., 2015; Mellough et al., 2015). Generation of iPSC-derived retinal cells from patients bearing *PRPF* mutations provides an excellent tool to study the cellular and molecular impacts of splicing deficiencies in these cells. In this study, we have used fibroblasts from four RP type 11 patients with variable clinical severity caused by two different *PRPF31* deletion mutations (c.1115_1125del11 and c.522_527del6&IVS6+1to+10del) to derive iPSCs. Patient-specific iPSC and age-matched controls were differentiated into RPE and three dimensional (3D) retinal organoids in order to elucidate disease mechanisms and to identify cell-type and patient-specific target genes affected by *PRPF31* mutations. CRISPR/Cas9 correction of a *PRPF31* mutation in cells derived from an RP11 patient with very severe RP, resulted in rescue of molecular and cellular phenotypes, thus providing proof-of-concept evidence for the effectiveness of *in situ* gene correction.

## Results

### Derivation of PRPF31-iPSC and their characterisation

Dermal skin fibroblasts from three related RP type 11 patients with a *PRPF31* c.1115_1125del11 heterozygous 11 bp deletion mutation **(Table S1)** with variable phenotypic expression of the disease, and one patient with severe RP with a *PRPF31* c.522_527+10del16 heterozygous 16 bp deletion mutation, were reprogrammed to iPSC using a non-integrative RNA-based Sendai virus **(Figure S1A).** Disease severity was determined according to fundus examination, visual field and visual acuity, and took account of the age at the time of examination **(Table S1).** Hereafter, all patients are labelled as RP11 accompanied by M (moderate), S (severe) and VS (very severe). Three unaffected controls are referred to as WT1, WT2 (Wilmes et al., 2017) and WT3 throughout this manuscript **(Table S1).** All RP11-iPSC harboured the mutation identified in fibroblast samples **(FigureS1B-E),** expressed pluripotency markers **(Figure S2A, S2B),** were free of transgenes **(Figure S2C),** genetically identical to parent fibroblasts **(Figure S2D)** and clear of any genomic abnormalities **(Figure S2E).** Both patient-specific and control iPSC were able to differentiate into cells belonging to all three germ layers *in vitro* **(Figure S3A)** and *in vivo* **(Figure S3B).**

### RP11-iPSC-derived-RPE display functional and ultrastructural abnormalities and loss of cell polarity

Control and RP11-iPSC were differentiated into RPE cells using the differentiation protocol shown in **Figure 1A, 1B.** RPE cells derived from unaffected control iPSC displayed the typical expression of the apical RPE marker Na^+^/K^+^-ATPase and the basolateral marker BEST1. A minority of RP11 RPE expressed both markers **(Figure 1C).** Immunostaining of control RPE showed polarised cells expressing collagen IV on the basal layer and MERTK in the apical layer **(Figure 1D).** In contrast, RPE monolayers derived from RP11 patients had reduced collagen IV and MERTK expression in the basal and apical layer **(Figure 1D).** Cytokine secretion assays revealed a significantly higher basal VEGF and apical PEDF expression in the severe and very severe RP11 patients when compared to the controls **(Figure 1E and F).** Collectively, these data indicate a loss of apical-basal polarity in patient iPSC-derived RPE cells.

**Figure 1.**
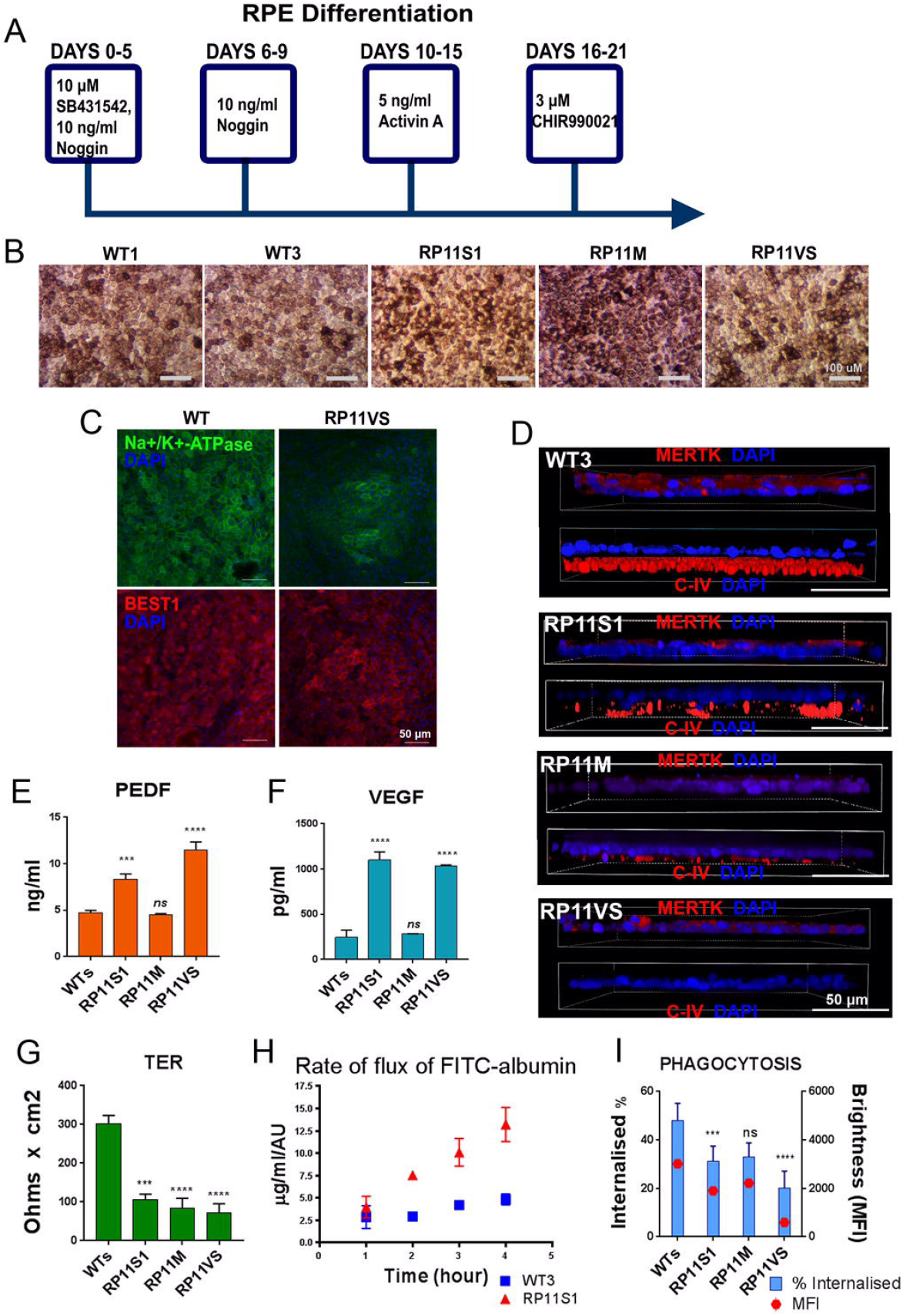
(see also figure S4). Differentiation of iPSC to RPE cells revealed polarity and functional defects. **(A)** Schematic of RPE differentiation timeline; **(B, C)** Bright field images and immunostaining for RPE markers; **(D)** Correct basolateral distribution of collagen IV (CIV) and MERTK in unaffected control (WT3) but not RP11 RPE cells; **(E, F)** ELISA assays for basal and apical secretion of VEGF and PEDF respectively in control and RP11 RPE cells. **(G, H)** Fluid transport and trans-epithelial resistance measurement revealed a significant difference between patient and RP11 RPE cells; (I) Reduced phagocytic capacity in RP11 RPE cells. E-I: Data shown as mean ± SEM, n=3. Statistical significance of pair-wise comparisons is indicated by n.s. not significant; *** p<0.001; **** p<0.0001 (Student’s paired t-test).

There was also a significant reduction in the ability of PRPF31-derived RPE cells to form a tight epithelial barrier as measured by trans-epithelial resistance assays **(Figure 1G)** and increased transport fluid from the basolateral to the apical side **(Figure 1H).** Furthermore, the ability to phagocytose rod outer segments was significantly reduced in RPE derived from the two patients with severe and very severe RP **(Figure 1I),** corroborating data previously obtained from knockdown of *PRPF31* in the ARPE19 cell-line (Farkas et al., 2014). Transmission electron microscopy (TEM) analysis carried out at week 21 and 43 of differentiation indicated the presence of typical RPE features including microvilli and melanosomes on the apical side of RPE derived from unaffected control iPSC **(Figure S4).** However, RP type 11 RPE displayed shorter and fewer microvilli, and contained large basal deposits underneath the RPE **(Figure S4)** that were not evident in controls.

### RP11-iPSC-derived photoreceptors display progressive degenerative features

RP11 and control iPSC were differentiated to 3D retinal organoids using the method published by Kuwahara et al. (2015) and shown in **Figure 2A.** Apical bright-phase neuroepithelium situated on the apical side of the retinal organoids was observed as early as day 20-24 of differentiation, and continued to grow in thickness throughout the differentiation process **(Figure 2B** and **Figure S5A).** By week 21, these 3D retinal organoids had a well-developed apical layer packed with photoreceptor precursors (marked by RECOVERIN expression) that contained connecting cilia (expressing ARL13B; **Figure 2C),** some of which had acquired a rod precursor phenotype as shown by NRL expression. Bipolar, ganglion and Müller glia cells were also present as evident from PKCα, HuC/D and CRALBP expression **(Figure 2C).** TEM of 3D retinal organoids at week 21 revealed the presence of outer limiting-like membrane (white arrows), inner segments (IS), connecting cilia (CC) and developing outer segments (OS) in photoreceptors residing in the apical layer of the 3D retinal organoids **(Figure 2D).** The same analysis repeated at week 43 of differentiation revealed the presence of asymmetric vacuoles (marked with black stars in **Figure 2E)** in the patient derived 3D retinal organoids that suggest “adaptive survival” in response to environmental or oxidative stress. Furthermore, apoptotic nuclei distinguished by electron-dense dark structures indicative of condensation of nuclear chromatin (indicated by white arrows in **Figure 2E)** were present in RP11 photoreceptors but not unaffected controls.

**Figure 2.**
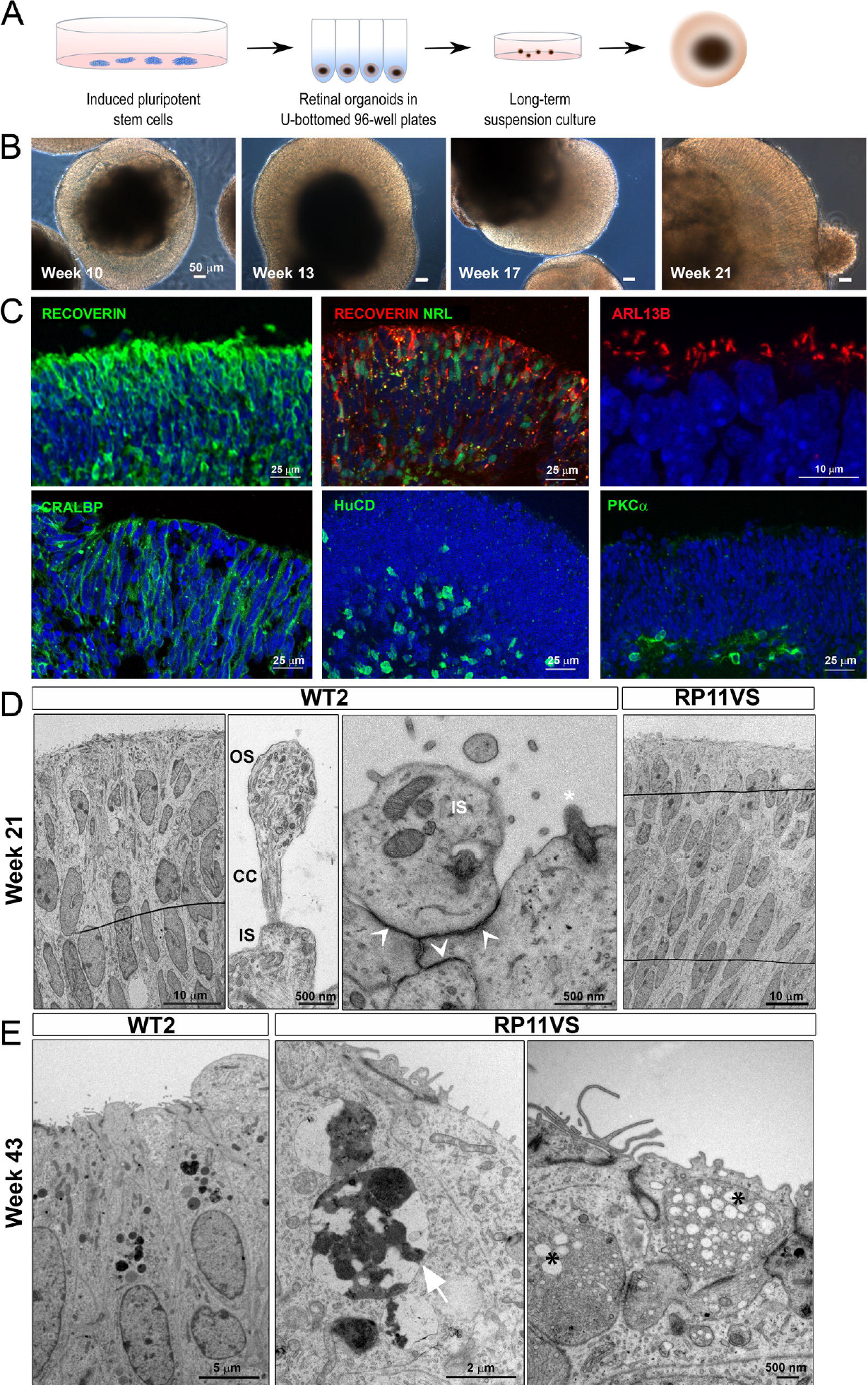
(see also Figure S5). Generation of 3D retinal organoids following long-term suspension culture. **(A)** Schematic representation of iPSC differentiation to 3D retinal organoids; **(B)** Bright field images showing development of retinal neuroepithelium over time; **(C)** Immunostaining of retinal organoids showing the expression of cell specific markers; **(D)** TEM revealed presence of outer limiting like membrane (white arrows), inner segments (IS), connecting cilia (CC), and developing outer segments (OS) in 3D retinal organoids after 21 weeks in culture; (E) After 43 weeks in culture, TEM showed that patient photoreceptors contained apoptotic nuclei with electron dense structures of condensed chromatin (white arrow) and stress vacuoles (black stars).

At week 21 of differentiation, the 3D retinae were flattened down on multi-electrode arrays (MEAs) with the presumed ganglion cell layer facing down on the electrodes in order to record action potentials generated by these cells. No differences between unaffected control- and RP11-retinal organoids were observed in response to 8-br-cGMP, a membrane permeable analogue of cGMP capable of opening the cationic channel associated with phototransduction in photoreceptor outer segments, triggering Na^+^ influx similar to the inward dark current **(Figure S5B, C** and **F).** Control retinal organoids responded to the addition of the neurotransmitter GABA with an increased firing rate, but this response was significantly reduced for the very severe RP11 patient **(Figure S5D, E** and **G).** GABA signalling emerges during very early development, and at that time it is depolarising and can induce spiking. It then becomes the main inhibitory neurotransmitter in the mature central nervous system. Hence, responses to GABA are indicative of emerging functional neural networks that are affected in RP11 patients.

### Impaired alternative splicing affecting genes involved in pre-mRNA splicing in patient-specific RPE and retinal organoids

To better understand the impact of *PRPF31* mutations, we performed semi-quantitative RT-PCR and western blot analysis of PRPF31 expression in primary fibroblasts, iPSC and iPSC-derived-RPE and 3D retinal organoids. The RT-PCR analysis across all the four cell-types indicated the presence of the nonsense mediated decay (NMD)-insensitive long mutant (LM) and NMD-sensitive short mutant (SM) transcript in cells derived from *PRPF31* patients only **(Figure 3A).** Quantitative measurements indicated a strongly diminished expression of *PRPF31* in iPSC-derived-RPE from PRPF31 patients **(Figure 3B).**

**Figure 3.**
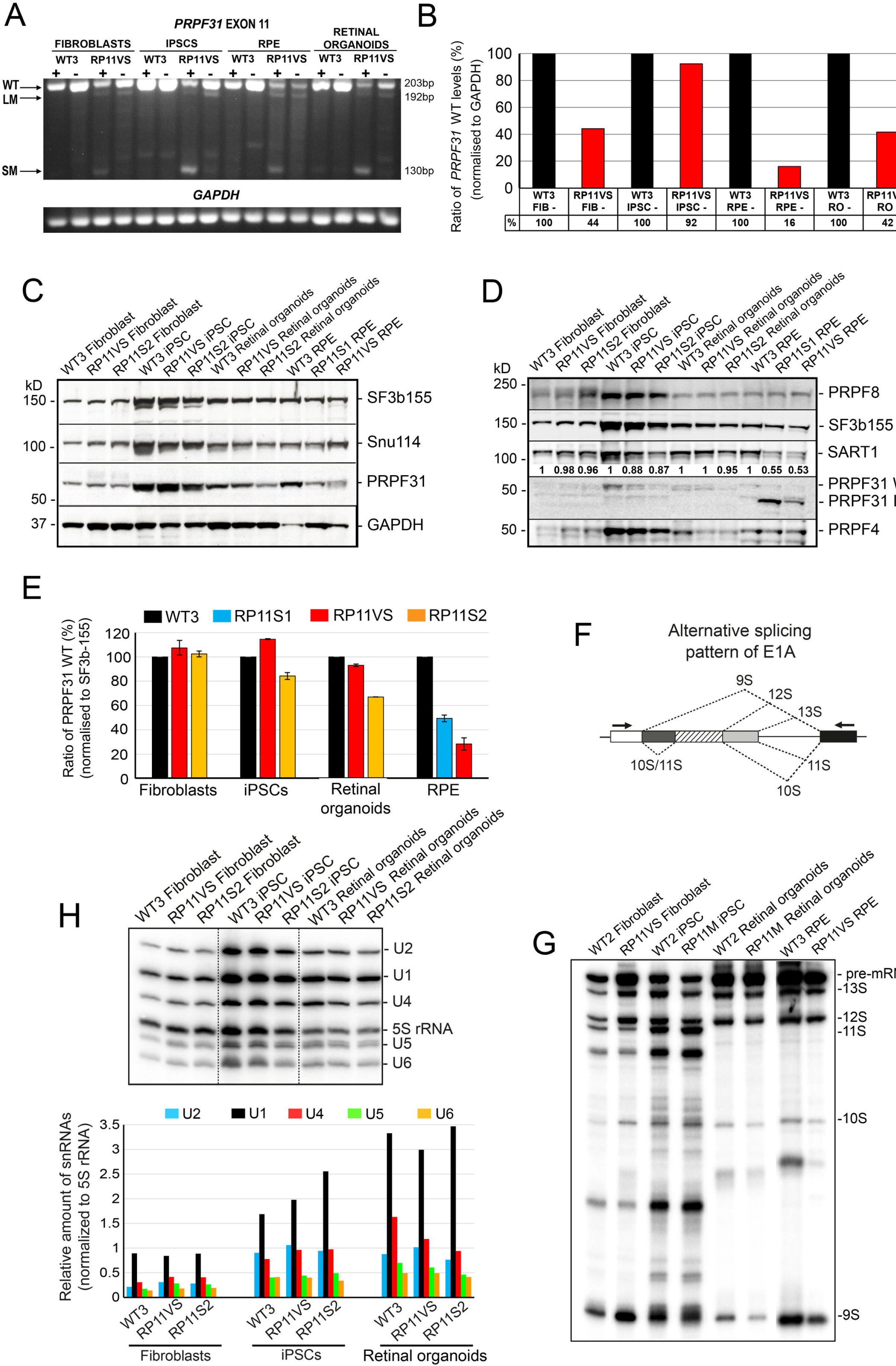
(see also Figure S6). PRPF31 expression in patient-specific cells and effects on pre-mRNA splicing. **(A)** Gel electrophoresis showing the presence of a long mutant transcript (LM) isoform for the exon 11 deletion in patient-specific cells. The short mutant (SM) isoform is present only upon inhibition of NMD with puromycin (indicated by +); **(B)** The bar graph shows wild type *PRPF31* mRNA in patient cells relative to controls from **(A).** Data are representative of at least 3 independent repeats, RO – retinal organoids; **(C-D)** Wild type PRPF31 is significantly reduced in patient RPE cells and less notably in 3D retinal organoids. The LM form and reduced SART1 is observed only in the patient RPE cells; **(E)** The bar graph shows wild type PRPF31 levels in patient cells relative to normal cells quantified from **(C); (F)** Patient RPE cells and 3D retinal organoids exhibit a notable defect in the alternative splicing of E1A mini-gene reporter. Schematic representation of alternative splice variants of the E1A reporter (F) and Denaturing PAGE and autoradiography using a phosphoimager (G); **(H)** Northern Blot analysis showing the level of snRNAs in various normal and patient cells. Total RNA was isolated from each sample and snRNA levels were analysed by denaturing PAGE followed by Northern blotting using probes against U1, U2, U4, U5, U6 and 5S rRNA (top). The levels of snRNAs were quantified and normalized to the amount of 5S rRNA (bottom).

Western blot analysis with an anti-PRPF31 C-terminus antibody indicated a significant down-regulation of PRPF31 protein in RP11 RPE, and to a lesser extent in the RP11 3D retinal organoids **(Figure 3C and E**). Interestingly, the presence of the PRPF31 LM (detected with a PRPF31 antibody against the N-terminus) was observed only in the RP11-RPE **(Figure 3D).** Furthermore, RP11 RPE also showed a substantial down-regulation of SART1, a U5 snRNP protein important for the formation of the pre-catalytic spliceosomal B complex, but no changes in the expression of the U5 protein PRPF8 or the U4/U6 protein PRPF4 **(Figure 3D).**

To understand whether splicing efficiency was altered in patient-specific cells, splicing assays (Caceres et al., 1994; Zerler et al., 1986a) were conducted by lentiviral transduction of an E1A mini-gene reporter with multiple 5’-splice sites, that can be alternatively spliced into at least five mRNAs (sizes 13S, 12S, 11S, 10S and 9S; **Figure 3F).** Both RP11 RPE and retinal organoids exhibited a notable defect in the alternative splicing of the E1A reporter, indicated by the accumulation of pre-mRNA and a decrease in 9S and 10S isoforms **(Figures 3F, G).** However, no defects were observed between unaffected control and patient-specific RP11 fibroblasts and iPSCs. Furthermore, there were no differences in tri-snRNP stability for RP11 and control iPSCs, as determined by sedimentation of nuclear extracts on density gradients followed by snRNA probing and immunoblotting **(Figure S6).** However, RP11 retinal organoids showed a lower expression of U4 snRNA **(Figure 3H;** red bars in bar graph) compared to controls, suggesting a likely reduced function of the U2-dependent spliceosome.

### RNA-seq analysis reveals alternative splicing of genes involved in pre-mRNA splicing, cells to substrate adherens junctions, ciliogenesis and DNA repair

To identify differences in transcription profile between RP11 patients and unaffected controls, RNA-seq analysis was carried out in fibroblasts, iPSC, RPE and retinal organoids **(Table S2).** Differentially expressed transcripts were identified in RP11 patients compared to unaffected controls for each cell-type by using DESeq2 with *p*_adj_<0.05 as the threshold **(Table S2).** Fewer differentially expressed genes were identified in iPSC (163) and RPE (59) samples when compared to fibroblasts (1395) and retinal organoids (1367). The genes with significantly changed expression between RP11 patients and unaffected controls in 3D retinal organoids **(Table S2)** showed enrichment of Gene Ontology (GO) categories related to actin cytoskeleton, ciliary membrane, primary cilium, photoreceptor inner and outer segments, axon terminals and phototransduction **(Table S3).** In fibroblasts, significantly changed genes were enriched in lysosome and endosomal process as well as focal adhesion, cell-substrate junctions and extracellular matrix organisation. There were no notable enriched pathways in iPSC and iPSC-derived-RPE differential gene expression datasets.

In view of the suggested impaired pre-mRNA splicing observed in RP11 RPE and retinal organoids **(Figure 3F, G),** and published reports of the role of PRPF31 in pre-mRNA splicing in photoreceptors (Yuan et al., 2005), we next identified target transcripts characterised by skipped exons, retained introns, alternative 5’ and 3’ splice sites, and mutually exclusive exons **(Table S4)** using rMATS software (Shen et al., 2014) across all four cell types (p_*adj*_< 0.05 and inclusion difference >5% as thresholds). This analysis **(Figure 4A)** revealed that RP11 RPE harboured the highest percentage of transcripts containing retained introns and alternative 3’ splice sites **(Table S4).** GO enrichment analysis revealed different biological processes being affected in each cell-type **(Figure 4B).** In fibroblasts, the transcripts with significantly different splicing between RP11 patients and unaffected controls were expressed from genes within the cilium formation categories (cilium assembly, cilium organisation, microtubule organising centre and centrosome; **Table S5).** This suggests that PRPF31 has a role in fibroblast ciliogenesis and corroborates our previously published data on reduced cilia length and incidence in RP11 fibroblasts (Wheway et al., 2015). GO enrichment analysis for cellular components **(Figure 4C)** identified microtubules, mid-body and cleavage furrow. These are all components that are important for ciliogenesis to occur after cell division (Bernabe-Rubio et al., 2016). In iPSC, biological component enrichment analysis identified DNA recombination and DNA double strand break repair, whereas the cellular component enrichment analysis identified transcripts involved in centrosome, centriole and microtubule organising centre **(Figures 4B, C).**

**Figure 4.**
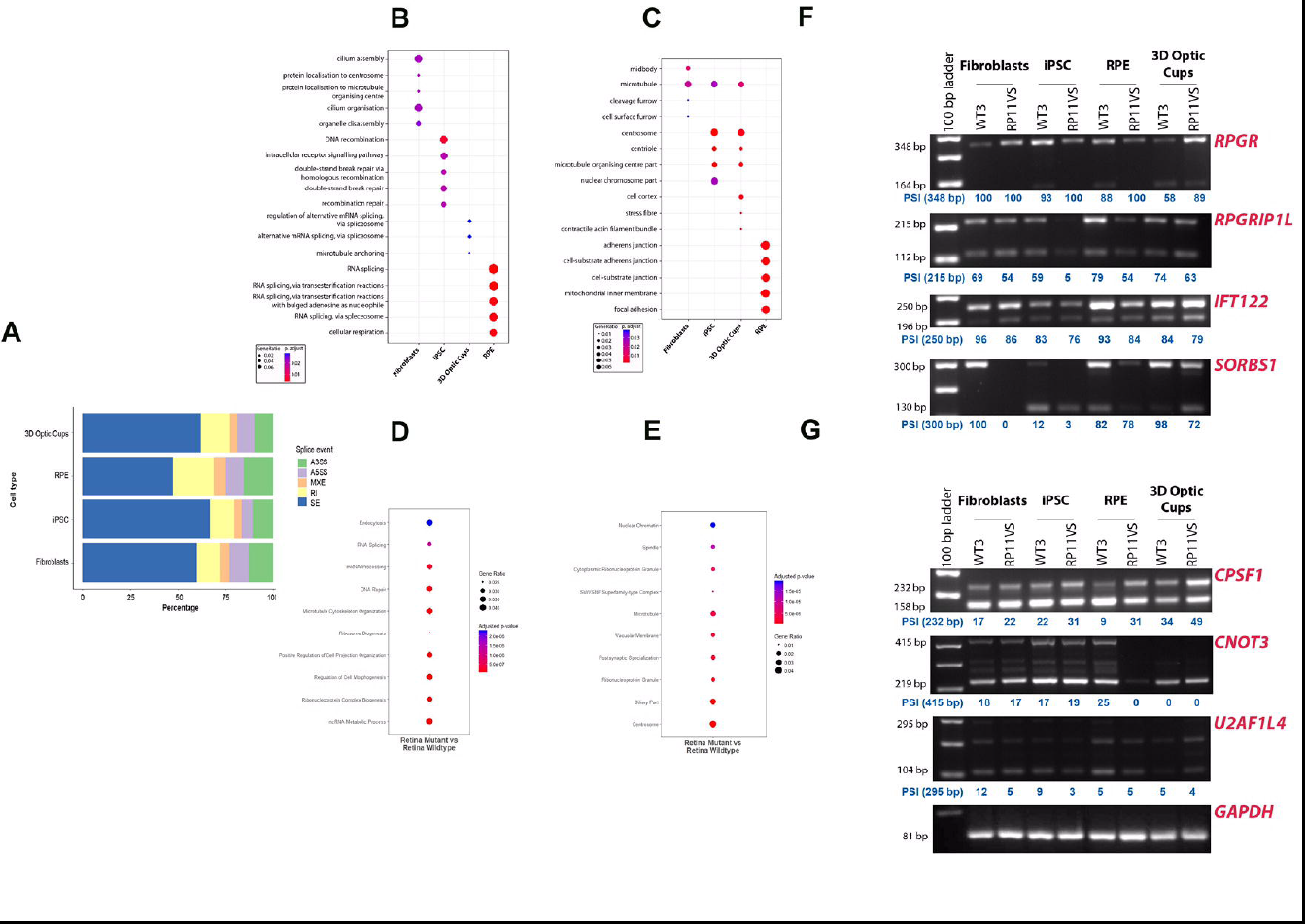
(see also Table S2-S5). RNA-seq analysis of alternative splicing in fibroblasts, iPSC, RPE and 3D retinal organoids and Prpf31 ^+/−^retina. **(A)** rMATS analysis showing that RP11 RPE have the highest percentage of transcripts containing retained introns (RI) and alternative 3’ splice sites (A3SS); **(B, C)** Gene Ontology enrichment analysis showing biological and cellular processes affected by alternative splicing respectively; **(D, E)** Gene Ontology enrichment analysis showing biological and cellular processes affected by alternative splicing respectively in the *Prpf31^+/−^* retinae; **(F, G)** Validation of RNA-seq findings by semi-quantitative RT-PCR, showing alternative splicing events in the key genes involved in pre-mRNA processing, cell adhesion and cilia. Blue numbers under lanes indicate the percentage spliced in (PSI) values for the major isoform. Loading control is for *GAPDH.*

Strikingly, in both RPE and retinal organoids, changes in alternatively spliced transcripts between RP11 patients and controls fell within the category of pre-mRNA splicing via the spliceosome **(Table S5** and **Figure 4B),** suggesting that *PRPF31* mutations lead to changes in alternative splicing of key components involved in the splicing process itself. Hence, this could result in exacerbation of splicing deficiencies and the unique cellular phenotypes observed in these mutant cells. The data obtained from human iPSC-derived-retinal organoids were also strongly corroborated by the results obtained from *Prpf31^+/−^* retinae (Graziotto et al., 2011; Farkas et al., 2014). Significant changes in alternatively spliced transcripts in mutant compared to wild-type controls occurred for the RNA splicing, mRNA processing and ribonucleoprotein complex biogenesis categories **(Figure 4D),** as well as microtubules, ciliary part and centrosomes **(Figure 4E).**

In human iPSC-derived-cells, the affected alternatively spliced transcripts are involved in various stages of mRNA processing and splicing including spliceosome assembly (e.g. *SF1, SART1, DDX5),* formation of the U4/U6 snRNP (*LSM2*), 3’-end processing of pre-mRNAs *(CPSF1, U2AF1L4),* association of U2 snRNP with pre-mRNA (*DDX39B, PTPB1)* and 5’-splice site selection (*LUCL7*). The cellular component analysis indicated that alternatively spliced transcripts were also involved in centriole and microtubule organisation in iPSC-derived-3D retinal organoids, and cells to substrate adherens junctions and focal adhesions in RPE cells **(Figure 4C, Table S5).** To validate these findings we performed RT-PCR experiments across all four cell types in RP11 and control samples **(Figure 4F, G)** to analyse key genes involved in cilia formation and/or outer segments of photoreceptors (*RPGR, RPGRIP1L, CNOT3),* actin filament organisation, centrosome and focal adhesion (*SORBS1*), intraflagellar transport (*IFT122*) and pre-mRNA 3’-end processing and splice site selection (*U2AF1L4* and *CPSF1).* The RP11 RPE showed the most significant change in alternative splicing of *CPSF1,* a protein that plays an important role in 3’-end processing of pre-mRNAs. The RP11 retinal organoids showed the highest degree of alternative splicing for *RPGR,* whilst RP11 RPE and iPSC displayed the highest degree of alternative splicing for *CNOT3* and *RPGRIP1L,* respectively **(Figure 4F, G).** Collectively these data suggest that *PRPF31* mutations result in the disruption of cell-type-specific regulation for alternative splicing targets within the retina.

GO cellular enrichment analysis indicated that the major processes affected in RPE due to alternative splicing are cell-substrate adherens junctions and focal adhesions **(Figure 4C).** These functions are intrinsically linked to the establishment of polarity in epithelial cells. The same analysis for RP11 retinal organoids revealed cilia and actin cytoskeleton components, thus linking aberrant splicing with cilia formation in photoreceptors.

### RP11 patient-specific RPE and photoreceptors show reduced cilia incidence and cilia length

To analyse the role of PRPF31 in cilia formation and function, we first analysed the expression of PRPF31 by immunocytochemistry using a snRNP-specific antibody (Y12, **Figure S7A)** and a cilia-specific antibody (ARL13B, **Figure S7B)** in combination with PRPF31 antibody in fibroblasts, iPSC, iPSC-derived-RPE and photoreceptors. Co-localisation with snRNP and ARL13B was observed across all patient cell types and controls, confirming PRPF31 localisation in splicing complexes and cilia, respectively.

We then measured cilia length and incidence in RP11 RPE cells using a combination of ARL13B and a centrosome marker (pericentrin) that is located at the base of the cilia **(Figure 5A).** Both cilia incidence and cilia length were significantly reduced in all patient-specific RPE cells when compared to controls **(Figure 5B).** TEM analysis indicated the presence of long cilia with clearly aligned microtubules in control RPE cells, whilst patient RPE cells displayed shorter, abnormal, bulbous cilia **(Figure 5C).** Structural defects in axonemal microtubules were confirmed by serial block-face scanning electron microscopy (SBFSEM, **Figure 5D).** Cilia analysis **(Figure 5E)** for photoreceptors indicated significantly lower cilia incidence in cells derived from RP11 patients **(Figure 5F).** TEM analysis indicated long cilia with clearly aligned microtubules in control photoreceptors and defective, bulbous cilia with misaligned microtubules in RP11 patients **(Figure 5G)** that was also confirmed by SBFSEM analysis **(Figure 5H).**

**Figure 5.**
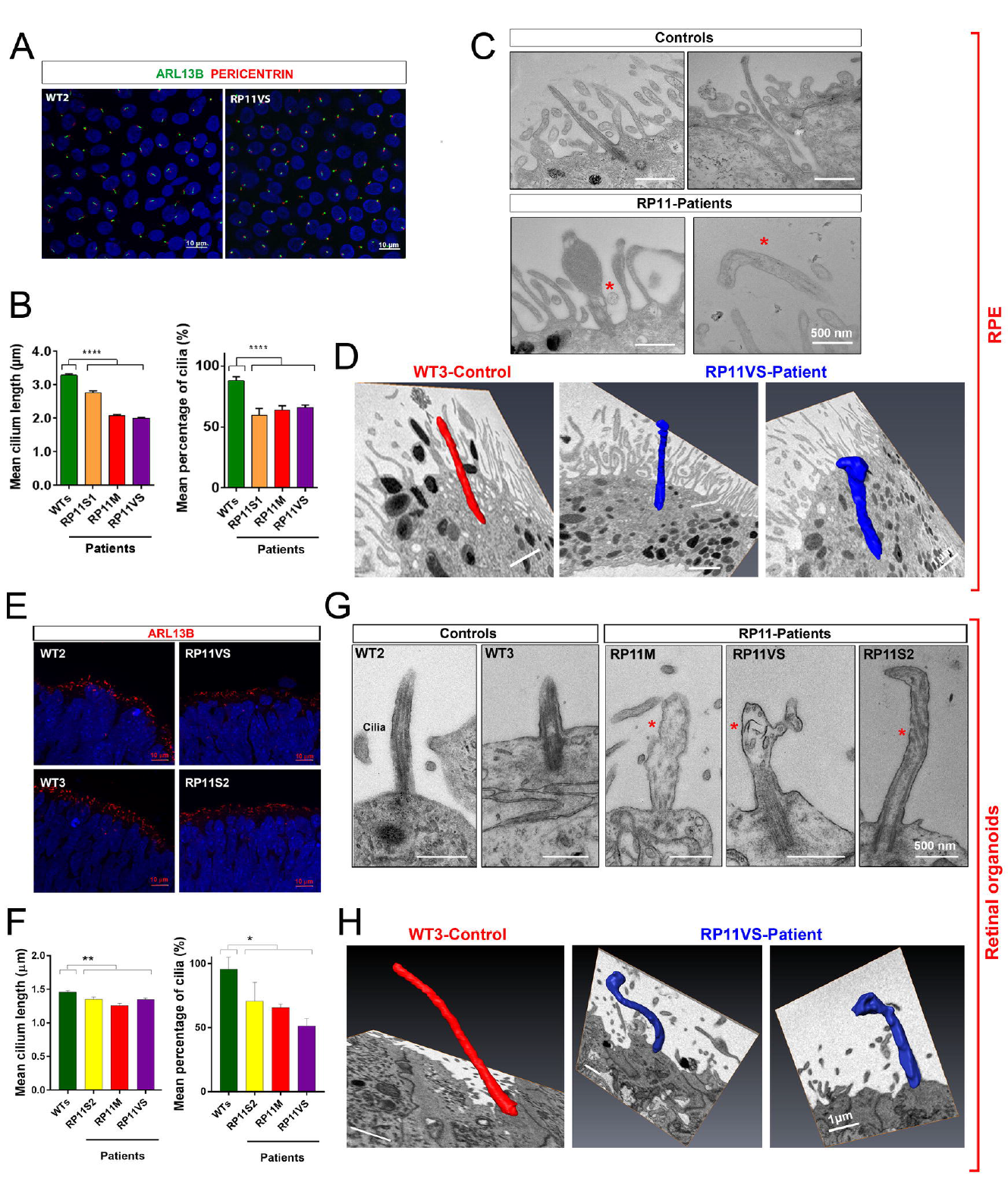
(see also S7). RP11 RPE cells and photoreceptors have defective ciliogenesis and cilia morphology. (A) Immunostaining of RPE with cilia markers ARL13B (green) and Pericentrin (red); **(B)** Quantification of cilia length and incidence showing significant reduction across both parameters in RP11 patients compared to the controls; **(C, D)** 2DTEM and 3D SBFSEM images showing shorter cilia in RP11 RPE cells, with abnormal bulbous morphology; **(E)** Immunostaining of photoreceptors with cilia marker ARL13B (red) at week 21; **(F)** Quantification of cilia length and frequency in photoreceptors showing significant reduction in RP11 patients compared to the controls; **(G, H)** 2D and 3D TEM images showing shorter cilia in patient-derived photoreceptors, with abnormal bulbous morphology. B,D, E, F: Data shown as mean ± SEM, n=3. Statistical significance of the indicated comparisons are indicated by n.s. not significant; *** p<0.001; **** p<0.0001 (One way ANOVA test with Dunnett’s post-hoc test correction for multiple testing).

To further confirm that human PRPF31 is a negative regulator of ciliogenesis, we performed siRNA knockdown in the human ciliated retinal pigment epithelial hTERT-RPE1 cell-line. This showed that knockdown of PRPF31 protein levels caused a significant decrease in cilia incidence **(Figure 6A).** SHH activity is known to require functional cilia, and we confirmed that *PRPF31* siRNA knockdown caused a dysregulated response to Smoothened agonist (SAG; **Figure 6B).** To investigate possible defects in cilia morphogenesis and structural organisation as a consequence of mis-splicing of ciliary genes, we used structured illumination microscopy (SIM) to resolve the detailed localization of proteins along the ciliary axoneme and at the transition zone (TZ). *PRPF31* knockdown caused significant mis-localisation of IFT88 to the ciliary tip **(Figure 6C).** The TZ proteins CC2D2A and RPGRIP1L were either entirely excluded from the TZ **(Figure 6D)** or mis-localised from the TZ into the ciliary axoneme **(Figure 6E)** following *PRPF31* knockdown. Similar results were obtained in RP11 RPE cells **(Figure 6F, G).**

**Figure 6.**
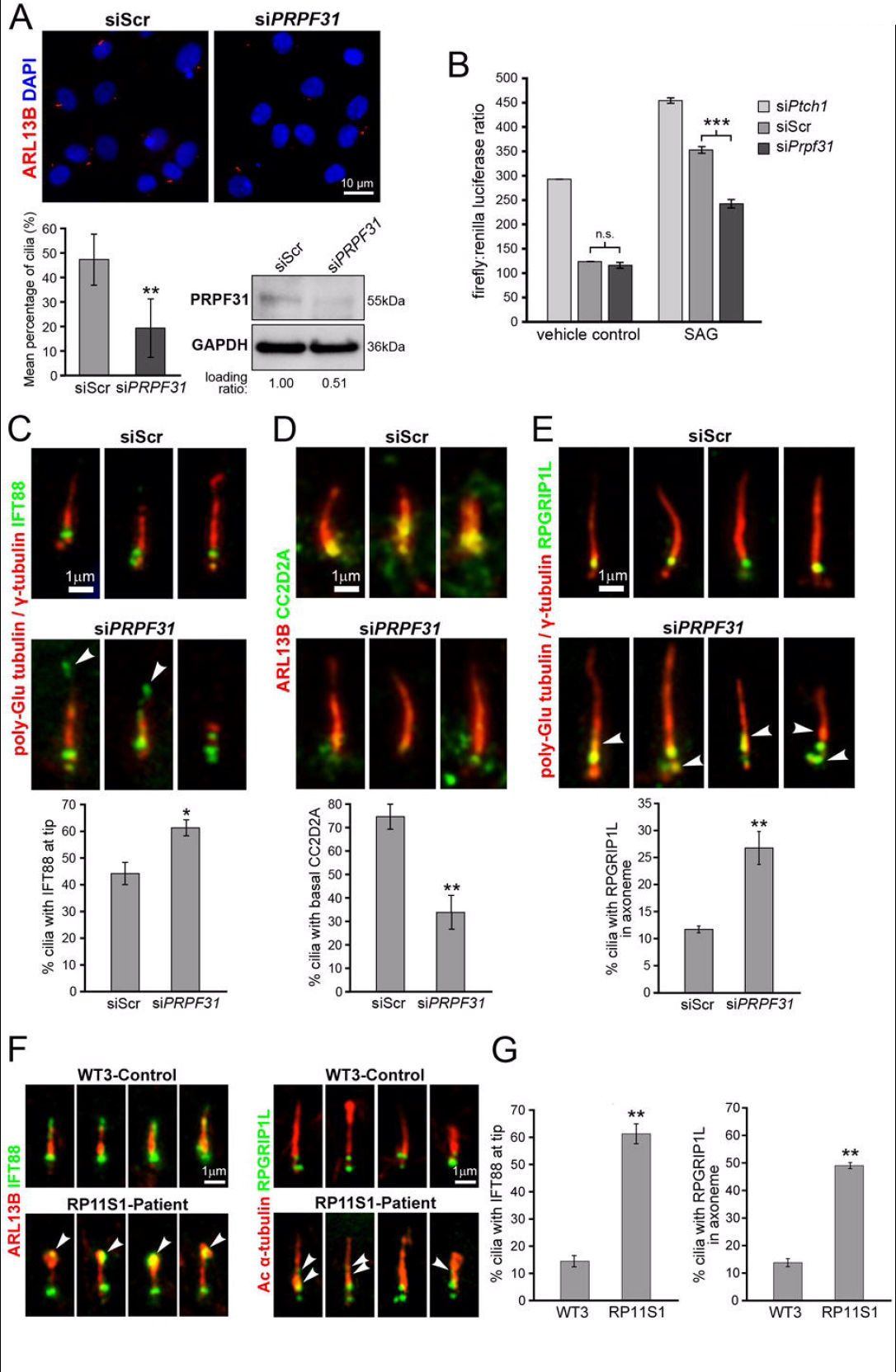
PRPF31 loss causes defects in cilia incidence and structural organisation. **(A)** *PRPF31* siRNA knockdown in human hTERT-RPE1 cells causes a significant decrease in cilia incidence (lower left) and PRPF31 protein levels (lower right) compared to scrambled negative control (siScr) siRNA; **(B)** Gli1 reporter assays of Shh activity measured in NIH3T3-GL cells following knockdown for *Ptch1* (positive control), scrambled negative control siRNA (siScr) and *Prpf31.* Cells were treated with either 100nM SAG or vehicle control for 48hr, as indicated. Assays results are expressed in arbitrary units of the ratio of firefly: *Renilla* luciferase activities; **(C)** Ciliary localization of IFT88 (green) in primary cilia of hTERT-RPE1 cells (visualized by staining for γ-tubulin and poly-glutamylated tubulin; red) showing mis-localization of IFT88 (arrowheads) at ciliary tips following PRPF31 knockdown. Bar graph quantitates the percentage of cilia with IFT88 at their tip. Scale bar = 1μm; **(D)** Visualization and quantitative analysis of the transition zone protein CC2D2A (green) and ARL13B (red); **(E)** Visualization and quantitative analysis of the transition zone (TZ) protein RPGRIP1L (green) and cilia (γ-tubulin and poly-glutamylated tubulin; red) showing mis-localization of RPGRIP1L from the TZ into the ciliary axoneme (arrowheads) following *PRPF31* knockdown. **(F-G)** Ciliary localization of IFT88 and RPGRIP1L (green) in RP11 RPE cells showing mis-localization of IFT88 (arrowheads) at ciliary tips and RPGRIP1L from the TZ into the ciliary axoneme (arrowheads). A-G: Data shown as mean ± SEM. Pair-wise statistical comparisons are indicated by braces: n.s. no significance.

### CRISPR/Cas9 mediated gene correction of *PRPF31* mutation restores key molecular and cellular defects in RP11 RPE and photoreceptor cells

To further validate the function of PRPF31 in retinal cells, CRISPR/Cas9 genome editing was used to correct the *PRPF31* c.1115_1125del11 genetic mutation in cells from the patient with the most severe clinical phenotype (RP11VS). For *in situ* gene correction, an ssODN template with wild-type *PRPF31* sequences was designed manually with 91bp homology arms on each side of the mutation region **(Table S6).** 200 iPSC clones were selected and tested as shown in **Figure 7A.** Candidates identified by PCR were sequenced to confirm gene-editing of *PRPF31* **(Figure 7B).** Quantitative RT-PCR analysis revealed increased expression of *PRPF31* in the CRISPR/Cas9 corrected clone **(Figure 7C)** when compared to uncorrected iPSC. The nine most likely genomic sites for targeting by CRISPR/Cas9 were predicted using the online tool CRISPR Design. These sites were amplified and sequenced, but no insertions or deletions were found at these predicted off-target sites **(Table S7).** CytoSNP analysis revealed that no genomic abnormalities had occurred during the *in situ* gene editing (data not shown). The CRISPR/Cas9 iPSC clone expressed pluripotency-associated markers Nanog and TRA-1-60, and gave rise to cells belonging to all three germ layers (data not shown).

**Figure 7.**
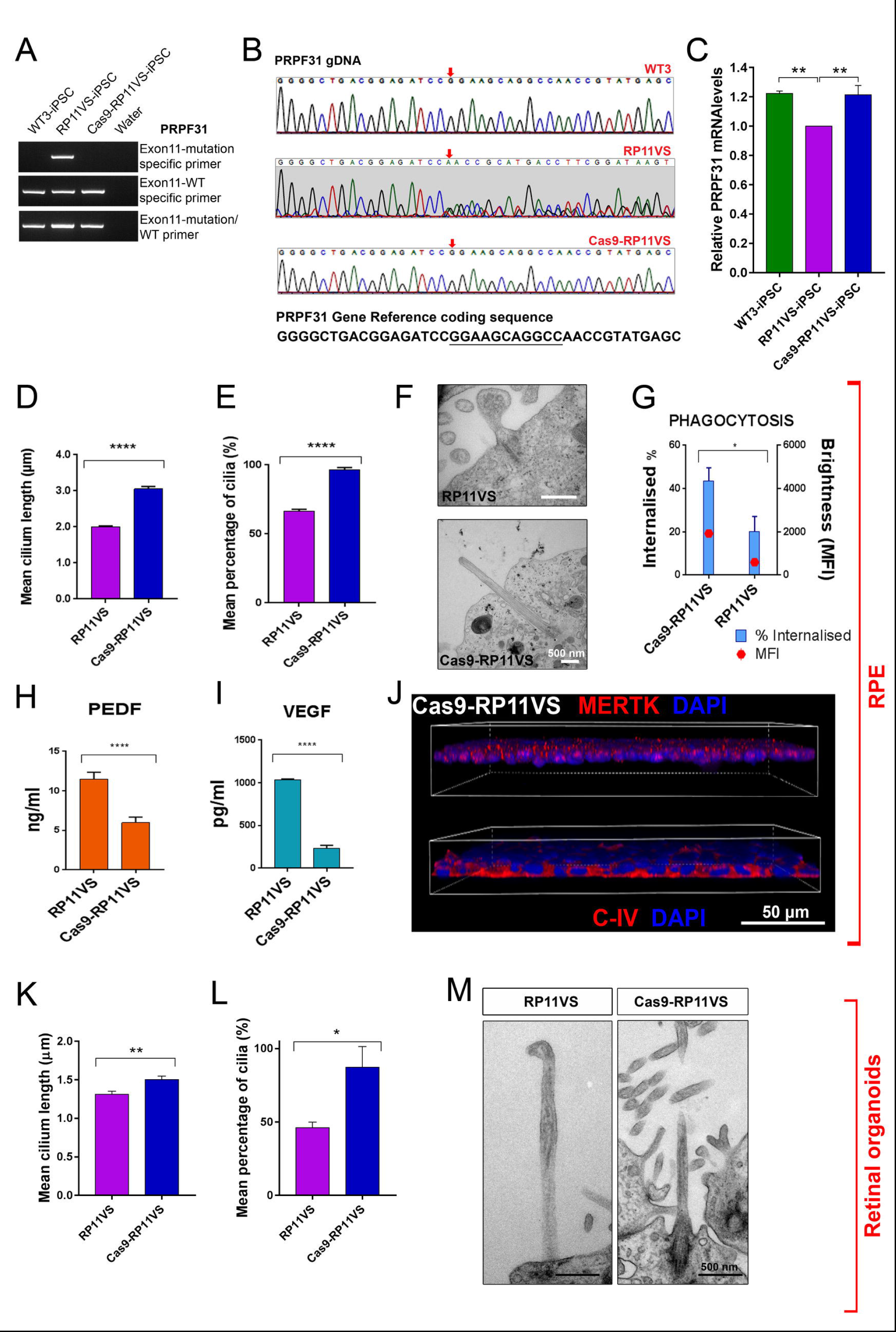
(see also Table S6 and S7). CRISPR/Cas9 mediated gene correction of *PRPF31* mutation in RP11 iPSC results in reversal of cellular and functional phenotypes in RPE cells and photoreceptors. (A, B, C) CRISPR/Cas9 correction of the *PRPF31* mutation in exon **11; (D, E)** Quantification of cilia length and incidence in PRPF31-edited RPE. **(F)** TEM analysis of *PRPF31-e*dited RPE cilia showing morphologically normal cilia; **(G)** Increased phagocytosis in *PRPF31-e*dited RPE. **(H, I, J)** Restoration of apical-basal polarity in *PRPF31-e*dited RPE. **(K, L)** Quantification of cilia length and frequency in PRPF31-edited photoreceptors; **(M)** TEM analysis of PRPF31-edited photoreceptor cilia showing morphologically normal cilia. C-E, G-I, K, L: Data shown as mean ± SEM, n=3. Statistical significance of pair-wise comparisons is indicated by n.s. not significant; * p<0.05; ** p<0.01; *** p<0.001; **** p<0.0001 (Student’s paired t-test).

The CRISPR/Cas9-corrected iPSC clone was differentiated to RPE and 3D retinal organoids alongside the uncorrected RP11 iPSC using our established protocols. A significant increase in cilia length and frequency was observed in both RPE **(Figure 7D, E)** and photoreceptors **(Figure 7K, L)** upon *in situ* gene editing of the *PRPF31* mutation. TEM analysis also revealed the presence of cilia with well-aligned microtubules that did not display the aberrant morphology observed in the RPll-derived retinal cells **(Figure 7F, M).** Importantly, flow cytometric analysis indicated the rescue of phagocytic capacity **(Figure 7G),** suggesting an improvement in RPE functional characteristics. Immunostaining and cytokine secretion assays revealed the restoration of cytokine secretion and basal collagen IV and apical MERTK expression, suggesting that RPE apical-basal polarity was restored **(Figure 7H, I, J).** These data indicated that *in situ* gene editing restored key cellular and functional phenotypes associated with RP type 11.

## Discussion

More than 90% of our genes undergo alternative splicing, generating the vast diversity of proteins in the human interactome (Wang et al., 2015). Mutations of spliceosome components would be expected to disrupt splicing function in all cells due to their ubiquitous requirement for splicing. However, mutations in spliceosome components produce distinct disease phenotypes such as skeletal dysplasia and retinal degeneration, raising tantalising questions as to why some tissues are more sensitive to splicing factor deficiencies. PRPFs fall under this category: they are ubiquitously expressed and involved in the formation of stable U4/U6.U5 tri-snRNPs and the spliceosomal B complex leading to spliceosome activation, yet human *PRPF* mutations result in retinal-specific phenotypes. Mouse models of RP type 11 have revealed late onset morphological and functional changes in RPE (Graziotto et al., 2011; Graziotto et al., 2008), whilst zebrafish models of RP11 develop rod photoreceptor degeneration (Yin et al., 2011) after injection of mutant human *PRPF31* mRNAs. Despite this progress, the disease mechanisms for PRPFs-related RP remain unclear, in particular whether RPE and/or photoreceptors are the primary affected tissue.

We took advantage of iPSC technology and generated RPE and 3D retinal organoids from four patients with two different *PRPF31* mutations and variable clinical severity. Our data shows a large number of ultrastructural, cellular and functional deficiencies in patient-specific RPE cells which include accumulation of basal deposits underneath the RPE, shorter and fewer microvilli and cilia, impaired apical-basal polarity, increased basolateral-apical fluid transport, decreased trans-epithelial resistance and decreased phagocytic capacity. Similar to RPE, photoreceptors display shorter, less frequent, defective cilia with misaligned microtubules suggestive of structural defects in axonemal microtubules. Collectively these data suggest that RPE is the most, but not the only, affected retinal cell type in *PRPF31-* mutated RP11 patients. Our data also show that patient-derived RPE cells not only harbour the lowest expression of remaining wild-type PRPF31 protein when compared to fibroblasts, iPSC and iPSC-derived-3D retinal organoids, but they also contain clearly detectable mutant PRPF31 proteins. These may act in a dominant-negative manner alongside PRPF31 wild-type proteins to compromise spliceosome function in a cell type specific fashion. In contrast, the PRPF31 mutant protein is not detected in patient-specific 3D retinal organoids, which suggests that any cellular and functional impairment in photoreceptors may be due to PRPF31 haploinsufficiency.

*In vitro* analysis across the four cell types suggested impaired global splicing in RPE and 3D retinal organoids, but not fibroblasts or iPSC, which was further corroborated by a lack of differences in U4/U6.U5 tri-snRNP complex formation between patient-specific and unaffected control iPSC. To better understand the difference between retinal cells (RPE and 3D retinal organoids) and other cell-types (fibroblasts and iPSC) we performed RNA-seq and identified transcripts with significantly altered splicing in comparisons between patient and unaffected control data-sets. Enrichment analyses for ontology showed that in RPE and 3D retinal organoids, but importantly, not fibroblasts or iPSC, the transcripts with the most significant change were in the pre-mRNA splicing category. This was corroborated by similar analysis of *Prpf31*^+/-^ mice that showed that alternatively spliced transcripts in retinae but not brain (data not shown) also fell within the pre-mRNA splicing category. Both our *in vitro* and *in vivo* data suggests that the exacerbation of splicing deficiencies in retinal cells leads to the restricted retinal phenotype in RP11 patients. Furthermore, the splicing deficiency is likely to have a serious deleterious effect on cellular phenotypes, since it is known that precise regulation of mRNA splicing in retina is essential for the development of primary cilia and photoreceptor outer segments (Murphy et al., 2016).

The structural integrity of connecting cilia is essential for the photoreceptor function (Parfitt et al., 2016). We observed shorter, defective and fewer cilia in patient-derived RPE and photoreceptor precursors. For the latter, we also observed changed alternative splicing of genes involved in cilia formation that further compromise pre-mRNA splicing patterns during the development of primary cilia in photoreceptors (Murphy et al., 2016; Parfitt et al., 2016). Nonetheless, splicing of ciliary genes was not as significantly affected in RPE, suggesting that cilia dysfunction in these cells may be due to other cellular phenotypes caused by altered splicing such as loss of apical-basal polarity which is tightly interlinked with cilia growth in epithelial and retinal cells (Krock and Perkins, 2014; Rodriguez-Boulan and Macara, 2014). The most affected cellular processes in RPE were the cell-to-substrate adherens junctions and focal adhesions. Disruption of these cellular phenotypes is the most likely cause for the reduced polarity and barrier function we observed in RPE cells derived from RP11 patients (Ma et al., 2013).

CRISPR/Cas9-mediated *in situ* gene editing has become a popular methodology for correcting mutations in iPSC prior to differentiation into cells of interest. We were able to correct the mutations in cells derived from the patient with the most severe clinical phenotype. This led to the reversal of key cellular and functional phenotypes in RPE and photoreceptors, without causing off-target effects, and thus providing proof-of-concept that this approach is effective. In summary, our data provide, for the first time, a mechanistic understanding of retinal-specific phenotypes in PRPF31-mutated RP type 11. Our studies highlight the advantages of iPSC-based disease modelling for identifying the affected retinal cell types and target genes, and for testing potential targeted therapies.

## Author Contributions

AB, LZ, VC, BB, SMJ, DD, AD, JC, RB, SM, MF, GH, KW, DH, KB, GC, CM, YD, SP, JAA, SA, YX, GW, KS, ES, LA: data acquisition and analysis, contributed to manuscript writing MM, CFI, DJE, SL, DHS, LA, ES: sample collection and data analysis SMJ, RRA, DHS, LA, EP, RL, SNG, CAJ, ML: study design, data analysis, fund raising and manuscript writing.

All authors approved the final version of the manuscript.

## Acknowledgments

The authors are grateful for financial support from Fight for Sight (1456/1457), ERC (CoG_614620 and 2012-ADG_20120314), RP Fighting Blindness (GR584), CiC (MC_PC_15030), SFB (860), Medical Research Council (MR/K011154/1, MR/M000532/1, MR/N005872/1), NIH (RO1 EY020902), and the Rosanne Silbermann Foundation and the European Community's Seventh Framework Programme FP7/2009 under grant agreement no. 241955 SYSCILIA. We would like to thank Newcastle University flow cytometry and microscopy core facilities, the MRC Medical Bioinformatics Centre at the University of Leeds and the Durham University computational biology facility for help with technical assays and RNA-seq analysis.

The authors declare no conflict of interest

**Figure Sl.Genotype of RP patients and corresponding iPSC. (A)** iPSC generation timeline; **(B)** Pedigree and phenotype of the patients with c.1115_1125del11 and mutation confirmation **(C); (D)** Pedigree and phenotype of the patient with c.522_527del6&IVS6+1to+10del and mutation confirmation **(E).**

**Figure S2 (see also Figure S1, S2 and Table S1). Generation and characterisation of patient and control iPSC. (A, B)** Bright field images and immunostaining for pluripotency-associated markers; **(C)** Verification of Sendai virus genome (*SEV*) and the transgenes (*KOSKLF4, C-MYC)* clearance by RT-PCR; **(D)** CytoSNP analysis confirming the genetic identity between iPSC and parent fibroblast and lack of genomic abnormalities **(E).**

**Figure S3 (related to Figure S2). iPSC were capable of spontaneous differentiation towards cells representative of three germ layers** *in vitro* **(A) and** *in vivo* **(B).**

**Figure S4 (related to Figure 1). Abnormal ultrastructural features of patient RPE cells.** TEM at 21 and 43 weeks in culture shows shorter and fewer microvilli in RP11 RPE cells and large basal deposits that were not observed in controls.

**Figure S5 (related to Figure 2). Spiking activity recorded from presumed retinal ganglion cells in 3D retinas derived from control (WT2) and diseased iPSC line (RP11VS) at week 21 of differentiation.** The activity was recorded with a 4096 channel MEA in contact with the tissue. **(A)** Brightfield images of developing retinal neuroepithelium over time; **(B)** Raster plot (top panel) and firing rate histogram (bottom panel) illustrate all the units in a WT2 sample showing an increase in firing in the presence of 8-br-cGMP. In the raster plot, each small vertical bar indicates the time stamp of an action potential. Each row represents a different RGC. The left half illustrates the activity in control conditions. The drug was puffed in the recording chamber (final concentration, 100 μM) at the time indicated by the green arrow and red line; **(C)** Same as in (B) but in a RP sample. The increase in activity is less conspicuous than in (B); **(D)** Same as in (B), but this time in the presence of the neurotransmitter GABA puffed as indicated by the green arrow and red line (final concentration 125 μM). Some cells exhibit a strong increase in firing when exposed to GABA in this WT2 sample; **(E)** Same as (D) but in a RP sample. The increase in activity is less conspicuous than in (D); **(F)** Box plot summarising cGMP-mediated increases in firing rate (expressed as % increase from control conditions) in WT2 and RP samples. The box plot shows the median (red line) and interquartile ranges with Tukey whiskers. There is no significant difference between WT2 and RP11VS (Mann Whitney test; N=20 for WT2 and N=37 for RP); **(G)** Same as (F) but for GABA. Here there is significantly less response in RP11VS samples than in WT2 (*, p<0.02; Mann Whitney test; N=24 for WT2 and N=26 for

**Figure S6 (related to Figure 3). Analysis of snRNP levels in RP11 and control iPSC. (A)** Northern blots of glycerol gradient fractions from normal and patient-specific iPSC nuclear extracts; **(B)** Top: Quantification of the U4 and U6 snRNP distributions across the gradient shown in A. Bottom: Ratios of U4/U6, U5 and U4/U6.U5 tri-snRNP in iPSC quantified from data in A; **(C)** Western blots of even-numbered glycerol gradient fractions of nuclear extracts from wild type and mutant PRPF31 iPSC. Migration positions of snRNPs are indicated below the gel and the detected proteins at the right.

**Figure S7 (related to Figure 6). Co-Localisation of PRPF31 with spliceosome component (Y12) and cilia (ARL13B) in fibroblasts, iPSC, RPE, and 3D retinal organoids.**

## STAR Methods

### CONTACT FOR REAGENT AND RESOURCE SHARING

Further information and requests for resources and reagents should be directed to and will be fulfilled by the Lead Contact, Majlinda Lako.

### EXPERIMENTAL MODEL AND SUBJECT DETAILS

#### Human Subjects

All samples used in this study were obtained with informed consent according to the protocols approved by Yorkshire & The Humber Research Ethics Committee (REC ref. no. 03/362). Further information on the patients and controls is provided in Method Details and in **Table S1.**

#### Culture of iPSC, RPE and 3D Retinal organoids

iPSC were generated and characterised as described in Method Details. Pluripotency was established by immunocytochemistry, flow cytometric analysis, and *in vivo* and *in vitro* differentiation into three embryonic germ layers. Cells were also tested for genetic identity and genomic abnormalities.

#### Cell culture

All cells were maintained in a humidified tissue culture incubator under the following conditions: 37°C, 5% CO_2_, 20% O_2_. Details of cell media composition can be found in Method Details.

### METHOD DETAILS

#### iPSC Generation

Three age-matched unaffected controls (WT1, WT2, WT3) and four RP11 dermal skin fibroblasts (RP11-1M, RP11-1S, RP11-2S and RP11-1VS, **Table S1)** were cultured with Advanced Dulbecco’s Modified Eagle Medium (Thermo-Fisher, Waltham, MA, USA) containing 10% FBS (Thermo Fisher Scientific), 1% Glutamax (Thermo Fisher Scientific) and 1% penicillin/streptomycin (Thermo Fisher Scientific) at 37°C and 5% CO_2_ in a humidified incubator. These fibroblasts were transduced at a density of 30,000 cells/cm^2^ using the CytoTune™-iPS 2.0 Reprogramming Kit (Life Technologies, A16517) following the manufacturer's instructions. iPSC colonies were established on inactivated primary mouse embryonic fibroblasts feeder layer and then adapted to the feeder-free system described below.

#### iPSC Culture

Human iPSCs were cultured on 6-well plates on Matrigel™ GFR (Corning, 354230) coated wells with mTeSR™l (StemCell Technologies, 05850) media supplemented with penicillin/streptomycin (Gibco, 15140). Cell culture medium was replaced on a daily basis. Cells were allowed to grow for 4-5 days prior to passaging or induction of differentiation. Passaging was carried out using Versene (EDTA 0.02%) (Lonza, BE17-771E) solution at 37°C for 3-5 minutes and cells were transferred to fresh matrigel plates in a 1:3 ratio. All cultures were maintained at 37°C, in a humidified environment, with 5% C0_2_. Cells were cryopreserved with freezing media containing 90% Fetal Bovine Serum (Gibco, 10270) and 10% Dimethyl Sulfoxide (Sigma, D2650).

#### Detection of Pluripotencv Markers by Immunocvtochemistrv

iPSC colonies were fixed in 4% poro-formaldehyde (Sigma, 47608) for 15 minutes at room temperature and permeabilised with 0.25% Triton-X-100 (Sigma, T8787) for 40 minutes. Blocking solution was applied (10% FBS + 1% Bovine Serum Albumin - Sigma, A3311) for 45 minutes at room temperature before proceeding with addition of anti-human SSEA4 conjugated with Alexa Fluor^®^ 555 (BD Biosciences) and anti-human OCT4 primary antibody (R&D). Secondary staining was performed with the antibody anti-goat IgG with FITC (Sigma) diluted in blocking solution, followed by nuclear counterstaining with DAPI (Partec, 05-5005). Colonies were imaged using a Bioscience Axiovert microscope in combination with the associated Carl Zeiss software, AxioVision. All antibody details are shown in **Table S6.**

#### Detection of Pluripotencv Markers by Flow Cytometric Analysis

iPSCs were treated with 0.02% EDTA (Lonza, BE17-711E) for 3 minutes at 37°C to dissociate the colonies. The suspension was collected in Phosphate Buffer Saline (PBS) and centrifuged for 3 minutes at 300xg. Supernatant was removed and replaced with PBS with 0.1% BSA containing TRA-1-60 conjugated FITC (Millipore) and NANOG conjugated with Alexa Fluor^®^ 647 (Cell Signaling). Samples were incubated in the dark at room temperature for 60 minutes on a shaker. Cells were washed with PBS and resuspended in FACS buffer (PBS with 2% FBS). A FACS Canto II flow cytometer was used to identify cell populations based on cell size (Side Scatter), cell granularity (Forward Scatter) and fluorescence. Results were analysed using the FACSDiva software. At least 10,000 cells were analysed for each sample.

#### *In Vitro* and *In Vivo* Three Germ Laver Differentiation Assays

iPSCs were detached from six well plates (20-30 colonies per well) using 1 ml of 1 μg/ml Collagenase type IV (Gibco 17104-019) and 0.5 μg/ml Dispase II (Gibco, 17105-041) solutions. The colony suspension was transferred to a 50 ml conical tube until the colonies settled in the bottom of the tube. The supernatant was carefully aspirated and 2 ml of differentiation media, containing DMEM-F12 (Gibco 11330), 20% FBS (Gibco, 10270), 1% Penicillin/Streptomycin (Gibco, 15140), 1% Non-Essential Amino acids (Gibco, 11140), was added per well. The colony suspension was then transferred to a 10 cm petri dish and media was changed every day. After 7 days, the embryoid bodies (EBs) were transferred to a gelatin coated 24 well plate or a chamber slide. After an additional 7 days colonies were fixed and stained with specific antibodies for the three germ layers using the 3-Germ Layer Immunocytochemistry Kit (Life Technologies, A25538). As a negative control, cells were stained only with secondary antibodies.

For the teratoma assay, iPSC colonies were dissociated with EDTA and 1 million cells were resuspended in a 200 μl solution of PBS (Gibco, 14190) + 2% FBS (Gibco, 10270). The samples were injected intraperitoneally in immunosuppressed mice at the Comparative Biology Centre, at the Medical School, Newcastle University. Each injection consisted of 0.5 million cells mixed with 100 μl Matrigel (BD, 354230). Following a period of 10 weeks, the mice were euthanized and the teratomae were excised, processed and sectioned according to standard procedures and stained for Weigert's haematoxylin, Masson's trichrome and Mayer's haematoxylin and eosin histological analysis. Sections (5-8 μm) were examined using brightfield microscopy and stained tissue photographed as appropriate.

#### hTERT-RPE1 Cell Culture

Human telomerase reverse transcriptase-transformed retinal pigment epithelium (hTERT-RPE1) were purchased from American Type Culture Collection (ATCC) at passage 9. The genomic status of the cell-lines were assessed by array CGH and karyotyping (May 2015). All cell-lines were tested every three months for mycoplasma. Cell-lines were maintained in DMEM/Ham's F12 medium (Gibco, 31331) supplemented with 10% Foetal Calf Serum (FCS) (Sigma-Aldrich, F7524), under standard conditions (37°C, 5% CO_2_). Cells were passaged at a split ratio of 1:8 twice a week, with low passages (<25) for both mIMCD3 and hTERT-RPE1 cells. hTERT-RPE1 cells were serum starved in OptiMEM (Gibco, 31985) for 48 hours to induce ciliogenesis. Human dermal fibroblasts were derived from skin biopsies and cultured in fibroblast growth medium (Genlantis Inc.) as described previously (Wheway et al., 2015).

#### RNA Isolation and Reverse Transcription

iPSC and iPSC-derived cell pellets were washed with PBS before being lysed with RNA Lysis buffer provided by the RNA extraction kit ReliaPrep™ RNA Cell Miniprep System (Promega, Z6010). The manufacturer’s instructions were followed, including a DNAse incubation step to the extracted RNA. The products were then passed through a column and resuspended in nuclease-free water. RNA was stored at -80°C or immediately used for cDNA synthesis. RNA was measured with a NanoDrop 2000 Spectrophotometer (Thermo Scientific) and 1 μg of extracted RNA was converted into cDNA using GoScriptTM Reverse Transcription System (Promega, A5000) following manufacturer’s instructions.

#### Reverse Transcription Polymerase Chain Reaction (RT-PCR)

For detection of any residual expression of the ectopically applied Yamanaka factors, RT-PCR was utilised in which the primers used were complementary to part of the SeV vector as well as the transgene. Oligonucleotides for the housekeeping gene (*GAPDH*) were used as a positive control. For the detection of mRNA transcripts as a result of c.1115_1125 del11 mutation, we have designed primers to detect wild type, long and short mutant transcripts of *PRPF31* gene. All primers are listed on **Table S6.** For the PCR reaction mixture, cDNA produced from 1 μg of RNA was amplified using the primers described on CytoTune™-iPS 2.0 Sendai Reprogramming Kit User Guide at the concentration of 10 μM each in addition to 10 μM dNTP mix, 5X Green GoTaq^®^ Reaction Buffer and GoTaq^®^ DNA Polymerase (5 μl) (Promega, M3175). The PCR consisted of a 35-cycle program of 95°C for 30 seconds, 55°C for 30 seconds followed by 72°C for 30 seconds and was carried out using a Mastercycler^®^ thermal cycler. Following the reaction, the samples were analysed using a 2% agarose gel electrophoresis mixed with GelRedTM Nucleic Acid Stain (Biotium, 41003). A 100 bp ladder was run against the samples.

#### Quantitative Real Time Polymerase Chain Reaction (qRT-PCR)

qRT-PCR was performed using the GoTaq™ qPCR Master (Promega) according to the manufacturer’s instructions. Each reaction contained 5 μlGoTaq qPCR Master Mix (Promega), 0.5 μl cDNA sample, nuclease-free water, and 0.6 μl primers (10 μM). All amplified produced ranged from 100 to 200 bp in size. The plates were run on an Applied Biosystems 7500 fast Real Time PCR machine. The cycling program consisted of a hot-start activation at 95°C for 5 minutes, followed by 45 cycles of denaturation at 95°C for 10 seconds, annealing/extension at 60°C for 30 seconds, and denaturation 95°C for 1 minute. Following amplification, a melt-curve analysis was performed from 65°C to 95°C with 0.5°C increments every 10 seconds. Each sample was run in triplicate, and the average quantification cycle (Cq) value was determined. Control reactions were run with water instead of template for each primer pair to check for primer-dimers and reagent contamination. Normalised gene expression values (against *GAPDH)* were obtained using the ΔΔCT method. All primer details are shown in **Table S6.**

#### Genomic DNA Extraction

Genomic DNA was extracted from the pelleted cultures of the iPSC and corresponding parental fibroblast cell lines using the QIAamp DNA Mini Kit (Qiagen, 56304) following manufacturer’s instructions.

#### Mutation Screening

10 ng of DNA from control and patients’ fibroblasts and iPSCs was amplified via standard PCR (40 cycles of 95°C for 30 seconds, 64°C for 30 seconds, 72°C for 30 seconds) using primers described in Dong et al. (2013) for the specific exons where the *PRPF31* mutations were located. The amplified products were purified using the QIAquick PCR Purification Kit (Qiagen, 28104) and quantified using the Qubit^®^ 2.0 Fluorometer. The sequencing files were analysed in the SeqScape v.2.5 software and forward and reverse sequences from both fibroblasts and iPSCs were aligned and compared with the *PRPF31* reference sequence gene (NG_009759.1) from Genbank to identify the *PRPF31* mutations. The consensus sequences from the forward and reverse sequences were then extracted from the software and pairwise aligned against the coding *PRPF31* sequence. Here the nucleotide designated as 1 commences at position 36 of GenBank accession number AL050369. All primer details are shown in **Table S6.**

#### SNP Array

DNA samples from the iPSCs and corresponding parental fibroblasts cell lines were analysed using the Infinium HumanCytoSNP-12 (Illumina, WG-320-2101) SNP array following manufacturer’s instructions. The results were analysed using the BlueFuse Multi 4.3 software (Illumina, San Diego, United States).

#### iPSC Differentiation to Retinal Pigment Epithelium (RPE)

iPSC colonies were grown to 80-95% confluency and all differentiation areas were removed. mTeSR™l media was replaced with 2 ml of differentiation medium [Advanced RPMI 1640, (12633, Gibco), GlutaMAX-1 (35050, Gibco), penicillin/streptomycin (Gibco, 15140) and B-27 (Gibco, 17504)] supplemented with 10 μM SB431542 (STEMCELL™, 72232) and 10 ng/μl Noggin (R&D Systems, 6057-NG-025) from days 0 to 5. From days 6-9 only 10 ng/μl Noggin (R&D Systems, 6057-NG-025) was added to the medium. From days 10-15, the medium was supplemented with 5 ng/μl Activin A (PeproTech, 120-14A) and from days 16-21 Activin A was replaced with 3 μM CHIR99021 (Sigma, SML1046). The cells were then fed every two days till first RPE patches appeared normally by week 4 of differentiation. RPE patches were mechanically picked and placed in TryPLE (10X) (Invitrogen, USA) for 30 minutes to dissociate the cells, agitated by gentle pipetting at 10, 20 and 30 minutes. Cells were sieved using a 100 μm cell strainer and re-plated at 4.5x 10^5^ cells per cm^2^ on 24 well plates or 0.33 cm^2^ PET hanging cell culture inserts (Merck Millipore; Billerica, United States) coated with PLO/laminin (50ng/μl) (Sigma-Aldrich, USA).

#### iPSC Differentiation to 3D Retinal organoids

The method for generating 3D retinal organoids from iPSC was based on a previously described protocol (Kuwahara et al., 2015) with minor modifications. Briefly, iPSCs were dissociated into single cells using Accutase (Gibco, A1110501). iPSCs were re-aggregated using low-cell adhesion 96-well plates with U-bottomed conical well (Lipidure^®^ COAT Plates, NOF Corp.) at a density of 12,000 cells/well in mTeSR1 media supplemented with ROCK inhibitor (Y-27632, Chemdea, CD0141, 20 μM). After 48 hours, the media was changed to differentiation medium (45% Iscove’s modified Dulbecco’s medium (Gibco,12440-053), 45% Hams F12 (Gibco,31765-029), 10% KSR (Gibco,10828-028), Glutamax (Gibco,35050-038), 1% chemically defined lipid concentrate (Thermo, 11905031), 450 μM monothioglycerol (Sigma, M6145), penicillin/streptomycin (Gibco, 15140-122). This was defined as day 0 of differentiation. BMP4 (55 ng/ml, R&D, 314-BP) was added to the differentiation medium on day 6. Half of the medium was exchanged every 3^rd^ day. On day 18, the aggregates were transferred from a 96-well plate to a low-attachment 6-well plate, and further cultured in suspension in the neural retinal differentiation (NR) medium containing DMEM/F12 (Gibco, 31330-038), 10% foetal bovine serum (Gibco,10270-106), 1% N2 supplement (Thermo, A1370701), 0.1 mM taurine (Sigma,T8691), 0.5 [iM retinoic acid (Sigma,R2625), 0.25 μg/ml Fungizone (Gibco,15290-02), penicillin/streptomycin (Gibco, 15140-122). The cells were maintained for up to 21 weeks, with media changes every 3-4 days.

#### APS-MEA Experiments

Recordings were performed on the BioCam4096 platform with BioChips 4096S+ (3Brain GmbH, Lanquart, Switzerland), integrating 4096 square microelectrodes. Organoids were transferred to 33°C artificial cerebrospinal fluid (aCSF) containing the following (in mM): 118 NaCl, 25 NaHCO_3_, 1 NaH_2_ PO_4_, 3 KCl, 1 MgCl_2_, 2 CaCl_2_, 10 glucose, and 0.5 l-Glutamine, equilibrated with 95% O2 and 5% CO2. Organoids were opened longitudinally and placed, with the presumed RGC layer facing down, onto the 4096 channel MEA, flattened with a translucent polyester membrane filter (Sterlitech Corp., Kent, WA, USA). The organoids were allowed to settle for at least 2 hours. To reliably extract spikes from the raw traces we used a quantile-based event detection (Muthmann et al., 2015) and single-unit spikes were sorted using an automated spike sorting method for dense, large-scale recordings (Hilgen et al., 2017b). Statistical significance and firing rate analyses were evaluated by using Prism (GraphPad, CA) and MATLAB (Mathworks, MA). Light stimuli were projected as described previously (Hilgen et al., 2017). Broad white light pulses (200 ms, 217 μW/cm^2^ irradiance, 1Hz) were presented for 5 min onto the organoids after recording of 5 min without pulsed light stimulation. The drugs cGMP (8-Bromoguanosine 3’,5’-cyclic monophosphate, Sigma-Aldrich, MO) and GABA (γ-Aminobutyric acid, TocrisBioscience, Bristol, UK) were puffed in the recording chamber (final concentrations, cGMP 100 μM, GABA 125μM) and 2 minutes before and after the puff were recorded.

#### CRISPR/Cas9 Correction of *PRPF31* Mutation in the RP11VS

Correction of *PRPF31* mutation in the RP11VS iPSCs was achieved by using the CRISPR/Cas9 system in combination with ssODNs as homologous templates covering the mutation site. The online design tool (http://tools.genome-engineering.org) was used to design the sgRNA sequences and predict off-targets. The sgRNA (see **Table S6),** which targets only mutant but not wild-type *PRPF31* sequences and is with low off-targets, was chosen. The sgRNA was cloned into the CRISPR/Cas9 vector (pSpCas9(BB)-2A-Puro) following the protocol from Ran et al. (2013). The ssODN template with wildtype *PRPF31* sequences was designed manually with 91bp homology arms on each side of the mutation region **(Table S6).** The sgRNA-CRISPR/Cas9 vector and ssODN were co-transfected into the RP11VS iPSCs by using Lipofectamine-3000 (Invitrogen) according to the manufacturer’s instructions. 24 hours after transfection, puromycin (0.2 μg/ml) was added for 2 days. 4-5 days after selection, the resistant iPSCs were dissociated into single cells using Accutase (Gibco, A1110501). 100,000 cells in mTeSR1 media supplemented with ROCK inhibitor (Y-27632, Chemdea, CD0141, 20 μM) were plated on a 10 cm Matrigel-coated dish. After 7 days, the colonies were picked and transferred to a cell culture 96 well plate. When the wells became confluent, iPSCs were split in two 24 well plates for further expansion and DNA isolation. Genomic DNA (gDNA) was isolated using QIAamp DNA Mini Kit (Qiagen, 56304). Subsequently, PCRs were performed with the primers including PRPF31-mutation specific primers; *PRPF31-VJ1* specific primers; PRPF31-mutation/WT primers **(Table S6).** The positive clones, which are negative for PRPF31-mutation specific primers and positive for *PRPF31-VJ1* specific primers were sequenced to confirm *in situ* gene-editing of *PRPF31.*

#### Off-Target Prediction and Capture Sequencing

sgRNA off-target sequences were predicted using the online design tool (http://tools.genome-engineering.org). Each sgRNA off target sequence was blasted against the human genome reference (https://blast.ncbi.nlm.nih.gov/Blast.cgi). Capture intervals were expanded by approximately 500 bp in both the 5′ and 3′ directions. Primers were designed in this region **(Table S6).** The PCR products were then sequenced to check the off-target effects of sgRNA.

#### Measurement of Trans-Epithelial Resistance (TER)

TER was performed using a Millicell ERS-2 Voltohmmeter (Millipore, MERS00002) by measuring the resistance of the blank transwell insert with PBS (Gibco, 14190) and the insert with RPE cells. The shorter and longer tips of the electrode were inserted in the transwell apical chamber and in the basolateral chamber, respectively. The resistance was measured twice in each transwell insert. The resistance reading of the blank was then subtracted from the resistance reading of the cells for each measurement. The results were multiplied by the membrane area value using the formula Unit area resistance = Resistance (Ω) x Effective Membrane Area (cm2), where the final value was given in ohms (Ω).

#### Rate of Flux of FITC-Albumin

RPE monolayers cultured on transwells were prepared for the flux assay by aspirating the spent media from the insert wells before sequential transfer of the inserts into three beakers of warm modified-Krebs buffer, which was also the transport buffer (composition in mM: NaCl, 140, KCl, 5.4, NaH_2_PO_4_, 0.3, KH_2_PO_4_, 0.3, MgSO_4_, 1.2, Glucose, 5, CaCl_2_, 2, HEPES, 10, TRIS base to pH 7.4). The monolayers were allowed to equilibrate for 30 minutes in transport buffer before initiation of flux. The temperature of the set up was kept at 37 °C. To initiate flux, the contents of the basolateral chambers were replaced with buffer containing 50 μg/ml FITC-albumin (Sigma-Aldrich, A9771). Sampling of the apical chamber was performed at defined intervals by transferring 50 μl of from the chamber into a clean 96-well plate. Buffer was replaced in the sampled chambers after sampling. The amount of FITC-albumin fluxed into the apical chamber were quantified by measuring fluorescence in the samples at excitation and emission wavelengths of 485 nm and 520 nm, respectively, using a FLUOstar Omega multi-plate reader (BMG Labtech, UK). Alongside the samples, known concentrations of FITC-albumin were used to create a standard curve. From this standard curve, the specific concentrations of FITC-albumin appearing in the apical chambers were calculated. To account for the difference in cell numbers between the monolayers, cell viability assay was performed at the end of the flux experiment. The monolayers were washed as described, and incubated with 100 μl of buffer containing 20 μl of MTS reagent (Promega, G3582) for 30 minutes. Thereafter, 50 μl of buffer was transferred to a clean 96-well plate and the absorbance taken at 490 nm. The absorbance was directly correlated to cell number, and thus the amount of FITC-albumin appearing in the apical chambers were normalised to the absorbance produced by the MTS reagent.

#### Transmission Electron Microscopy (TEM)

RPE and 3D optic cup samples were fixed with 2% gluteraldehyde in 0.1M sodium cacodylate buffer and sent to the Transmission Electron Microscopy facilities at Newcastle University, where samples were fixed with 2% glutaraldehyde in 0.1M sodium cacodylate buffer, post-fixed in 1% osmium tetroxide, dehydrated in gradient acetone and emebedded in epoxy resin. Ultrathin sections (70nm) were picked up on copper grids, stained with uranyl acetate and lead citrate and imaged using a Philips CM100 transmission electron microscope with high resolution digital image capture.

#### Serial Block Face SEM (SBFSEM)

Cells were fixed overnight in 2% glutaraldehyde in 0.1M sodium cacodylate buffer. Once fixed, the samples were processed using the heavy metal staining protocol adapted from Deerinck et al. (2010). Briefly, samples were incubated in a series of heavy metal solutions - 3% potassium ferrocyanide in 2% osmium tetroxide, 10% thiocarbohydrazide, 2% osmium tetroxide again, 1% uranyl acetate overnight, and finally lead aspartate solution. Between each step the samples were rinsed thoroughly in several changes of deionised water. Samples were dehydrated through a graded series of acetone and then impregnated with increasing concentrations of Taab 812 hard resin, with several changes of 100% resin. The samples were embedded in 100% resin and left to polymerise at 60°C for a minimum of 36 hours. The resin blocks were trimmed to approximately 0.75 mm by 0.5 mm and glued onto an aluminium pin. In order to reduce sample charging within the SEM, the block was painted with silver glue and sputter-coated with a 5nm layer of gold. The pin was placed into a Zeiss Sigma SEM incorporating the Gatan 3view system, which allows sectioning of the block *in situ* and the collection of a series of images in the z-direction. Multiple regions of interest were imaged at x2K magnification, 3000 x 1500 pixel scan, which gave a pixel resolution of approximately 15 nm. Section thickness was 50 nm in the z-direction. In the resulting z-stacks, cilia were identified and segmented manually using Microscopy Image Browser (MIB, University of Helsinki). The segmentations were imported into Amira (FEI) for construction of the 3D models.

#### Phagocytosis Assay

Rod Photoreceptor Outer Segments (POS) from Bovine (InVisionBioResources, 98740) were centrifuged at 2600xg for 4 minutes and the pellet was resuspended in 100 μl of Advanced RPMI (AdRPMI) 1640 medium (12633, Gibco). The POS were incubated with 0.4 mg/ml FITC for 1 hour at room temperature and agitated in the dark. POS were centrifuged at 2600xg for 4 minutes and washed three times with PBS (Gibco, 14190). Then, POS were resuspended in AdRPMI 1640 (12633, Gibco) + B-27 Supplement (Gibco, 17504) + 10% Fetal Bovine Serum (FBS) (Gibco, 10270) and the staining was confirmed under a Bioscience Axiovert microscope. RPE cells were treated with 1 × 10^6^ POS-FITC per cm^2^ and incubated for 4 hours at 37°C. RPE controls cells were also treated with the same number of non-stained POS and incubated for the same time. Cells were rinsed with PBS supplemented with calcium and were detached from the wells using 200 μl of Trypsin for 5-8 minutes. Trypsin was neutralised by the addition of 500 μl of AdRPMI 1640 medium + 10% FBS and POS were centrifuged at 300xg. Cell pellets were resuspended in FACS buffer (PBS with 2% FBS) and transferred to FACS tubes. 5 mM DRAQ5 (Biostatus, DR50200; 1:2500) was used to distinguish cells from debris and outer segments. Cells were washed with 0.2% Trypan Blue Solution (Sigma, T8154) to quench fluorescence from bound POS, washed with PBS and suspended in FACS buffer. Samples were analysed immediately on a LSRII flow cytometer and 10,000 events were collected per sample. Results were analysed using FacsDiva software.

#### RPE Cytokine Secretion Studies

Medium from basal and apical chambers of transwell inserts were collected from RPE cells of healthy controls and patients. The levels of PEDF and VEGF secretion were measured by using human PEDF-ELISA Kit (Cusabio, CSB-E08818h) and human VEGF-ELISA Kit (Life technologies KHG0111) according to manufacturer’s instructions.

#### RPE Characterisation by Immunocvtochemistrv

Cells were fixed in 4% Formaldehyde (Sigma, 47608) for 15 minutes at room temperature and permeabilised with 0.25% Triton-X-100 (Sigma, T8787) for 15 minutes, followed by treatment with blocking solution (3% BSA in PBS, Sigma, A3311) for 30 minutes at room temperature. Cells were treated with primary antibodies Anti-Bestrophin (Abcam, 1:300), Anti-Sodium Potassium ATPase (Alexa Fluor^®^ 488 conjugate) (Abcam, 1:50), Pericentrin (Abcam, 1:500), MERTK (Bethyl, 1:200), ARL13B (Proteintech, 1:500), Collagen IV (Abcam, 1:200), PRPF31 (Abnova, 1:500) and SNRPB Monoclonal Antibody (Y12) (Thermo, 1|:500), overnight at 4°C, and with secondary antibodies anti-rabbit FITC (Sigma, 1:500) or anti mouse FITC (Jackson Immuno Research, 1:500) and anti-mouse Cy3 (Jackson Immuno Research, 1:500) or anti-rabbit Cy3 (Jackson Immuno Research1:500) diluted in PBS for 1 hour at room temperature. Washes with PBS were carried out between and after treatments. Finally, cells were treated with the nuclear stain-DAPI (Partec, 05-5005), and imaged using a Nikon A1R Confocal microscope in combination with the associated NIS Elements software. All antibody details are shown in **Table S6.**

#### Immunofluorescence and Confocal Microscopy of hTERT-RPEl and RPE Cells

Immunofluorescence staining was performed as described previously (Dawe et al., 2009). hTERT-RPE1 cells were cultured as described above. Coverslips were seeded with 10^5^ cells and serum starved in OptiMEM for 48 hours after transfection to induce ciliogenesis. 25 pmoles of siRNA was reverse-transfected using Lipofectamine RNAiMAx (Invitrogen). Ciliated cells were fixed in ice-cold methanol for 5 minutes at -20°C, treated with 0.05% Triton X-100 in PBS for 5 minutes, and blocked in 1% non-fat milk in PBS for 30 minutes. Fixed cells were stained for 90 minutes with appropriate primary antibodies. Coverslips were then washed in PBS and stained for 1 hour with appropriate Alexa Fluor-conjugated secondary antibodies (Life Technologies). Coverslips were washed again with PBS before mounting in Prolong Gold anti-fade mountant (Molecular Probes).mIMCD3 cells were seeded at 2.5x10^5^ cells/well on sterile glass coverslips in six-well plates and fixed in ice-cold methanol. Cells were blocked in 1% non-fat milk in PBS for 30 minutes. Images were obtained using a Zeiss ApoTome structured illumination microscope (SIM), equipped with a 63x objective oil lens, or a Nikon A1R confocal microscope with X100 oil objective lens controlled by NIS-Elements AR 4.20.01 (Nikon) software. Optical sections were generated through structured processing using Axiovision 4.3 (Zeiss) or NIS-Elements AR 4.20.01 (Nikon) software. Images were analysed using Adobe Photoshop CS and FIJI software. Images were assembled with Adobe Illustrator CS. All antibody details are shown in **Table S6.**

#### 3D Optic Cup Characterisation by Immunocytochemistry

iPSC-derived 3D retinal organoids were fixed in 4% paraformaldehyde at room temperature for 20 minutes. Post-fixation 3D retinal organoids were incubated with 30% sucrose in PBS, and then frozen and cryosectioned. The frozen sections were stained for a panel of retinal specific antibodies. Antibodies against the following proteins were used at the indicated dilutions: RECOVERIN (Millipore, 1:800), NRL (Santa Cruz, 1:800), CRALBP (Abcam, 1:500), HuC/D (Invitrogen, 1:500), PKCα (BD Pharmingen, 1:500), ARL13B (Proteintech, 1:500), PRPF31 (Abnova, 1:500), SNRPB Monoclonal Antibody (Y12) (Thermo, 1:500). The following secondary antibodies were used: Anti-Mouse-IgG-FITC (Jackson Immuno Research, 1:500), Anti-Mouse-IgG-Cy3 (Jackson Immuno Research, 1:500), Anti-Rabbit-IgG-Cy3 (Jackson Immuno Research, 1:500), Anti-Goat-IgG-FITC (Jackson Immuno Research, 1:500). Nuclei were labelled with Blue-DAPI (Thermo). All antibody details are shown in **Table S6.**

#### Cilia Length and Frequency Measurements in RPE Cells

The length of cilia was measured by a 3D method using Imaris 8.3 Software (Bitplane Inc). Immunocytochemistry Z-stacks images of RPE samples from patients and controls were uploaded into Imaris. Surfaces were created to cover all the length of the cilia across the bottom and top of the stacks. The values were given in μm and a minimum of 150 cilia were independently measured for each sample. The cilia incidence was calculated by counting the number of cilia per cell in each image. Z-stacks were uploaded into Imaris software and spots were created for the blue (DAPI) and green (FITC) channels to cover all nuclei and cilia of cells. The percentage of cilia spots per nuclei spots was calculated and a minimum of 300 cells were counted per sample.

#### Cilia Length and Frequency Measurements in 3D Retinal organoids

Cryosections from 3D retinal organoids were stained with the ARL13B antibody (Proteintech). The images were obtained using Carl Zeiss laser-scanning microscope and Zen software. Maximum intensity projections of Z-stacks were used for the analysis. The measurements were performed in MATLAB. The workflow of the method is briefly explained as follows. CZI image files are first imported into MATLAB workspace using the ‘bfopen’ function written by Bio-Formats (Linkert et al., 2010). This allowed us to access channels individually as grayscale images. Hysteresis thresholding is used to segment the cells and filter out noise from the images (Zimmer et al., 2002). The concept of this method is to use dual thresholding values, such that all the pixels with intensity values above the upper threshold value are first marked as cell pixels, any neighbourhood pixel above the lower threshold that is connected to each of the first pass pixels are also classified as cell pixels. This produces segmentation with fewer isolated points, giving a better result than a simple high-pass thresholding. Segmented objects, such as noise, those sizes smaller than a user selected value are then removed. Finally, a watershed-based method is applied to the binary image to split clustered cells, details of this method can be found in Wang et al. (2015). Because all the pixels of each cell are connected and represented as a single region, an image region property measuring function (regionprops) in MATLAB was used to extract the information of each cell, such as, size, average intensity value and length. The cilia length reported was the average length of all cilia in the image. The frequency of ciliated cells was calculated as cilia numbers/total cell numbers (DAPI) in the same region X 100%.

#### RNA Sequencing

Total RNA was extracted from tissue using TRIzol (ThermoFisher Scientific Inc). RNA samples were treated with a TURBO DNA-free™ Kit (Ambion Inc.) using conditions recommended by the manufacturers, and then cleaned with a RNA Clean & Concentrator™-5 spin column (Zymo Research Corp.) RNA was tested for quality and yield using a NanoDrop 1000 spectrophotometer and an Agilent 2100 Bioanalyzer. To minimize bubble PCR artefacts, we used 100 μg of purified total RNA in library preparation, following the “TruSeq” Illumina protocol. In brief, RNA was polyA-selected, chemically fragmented to about 200 nt in size, and cDNA synthesized using random hexamer primers. Each individual library received a unique Illumina barcode. RNA-seq was performed on an Illumina HiSeq 2000 or HiSeq2500 instrument with six or eight libraries multiplexed per flow-cell lane using 100 bp paired-end reads. This resulted in an average of 250 million reads per lane, with an average of 40 million reads per sample. Raw reads were aligned to the human *(Homo sapiens)* full genome (GRCm38, UCSC mm10) using STAR, a splice-aware aligner (Dobin et al., 2013). GTF transcript annotation files were downloaded from Ensembl. Transcripts were assembled using STAR, followed by estimates of raw gene counts using HTSeq (Anders et al. 2014). Differential gene expression was analysed using DESeq2 (Love et al., 2014) with statistical significance expressed as a p-value adjusted for a false discovery rate of 0.01 using Benjamini-Hochberg correction for multiple-testing.

Alternate splicing analysis was then carried out using rMATS (Shen et al., 2014). For each comparison being made we used the sorted BAM files produced by STAR to run rMATS using default unpaired settings. Reported splicing changes were considered significant if they had a p-value less than 0.05 and a change in inclusion level difference of more than 5%. GO Enrichment Analysis was carried out on the genes found to have significant splicing changes via clusterProfiler (Yu et al., 2012). Multiple testing corrections were carried out using the Benjamini-Hochberg method with an adjusted p-value of less than 0.05 denoting significantly enriched gene ontology.

#### Production of Lentiviral Particles and Transduction of Target Cells

The minigene reporter encoding the adenovirus E1A transcript was subcloned from pMTE1A plasmid (Caceres et al., 1994; Zerler et al., 1986b) into the *Pme*I site of the pWPI lentiviral vector (Addgene; Trono lab). Lentiviral particles were produced in HEK293T cells grown in DMEM medium with 10% FBS. The cells were transfected with pWPI-E1A and the packaging plasmids psPAX2 and pMD2.G (Addgene) using PEIpro transfection reagent (Polyplus transfection). After 54 hrs, the medium containing lentiviral particles was centrifuged at 1,000 g for 5 min and cleared using a 0.45 μm filter. The lentivirus was concentrated using Amicon Ultra 100 kD MWCO centrifugal filter units (Millipore) and aliquots were stored at - 80 °C. For lentiviral transduction, cells were seeded in 6-well plates with 2 ml medium and infected with the concentrated lentivirus in the presence of 8 μg/ml polybrene (Sigma). After 24 hrs the culture medium was refreshed, and 36 hrs later cells were washed with PBS and harvested.

#### E1A Alternative Splicing Assays

Total RNA was extracted from cells transduced with the E1A lentivirus using an RNA extraction kit (Macherey Nagel). E1A alternative splicing was analysed by RT-PCR with 1 μg of the total RNA sample using the High-Capacity cDNA Reverse Transcription Kit (Applied Biosystems) and GoTaq DNA polymerase (Promega). PCR was performed with the 50-end radiolabelled exon 1 forward primer (5-GTTTTCTCCTCCGAGCCGCTCCGA) and the exon 2 reverse primer (5-CTCAGGCTCAGGTTCAGACACAGG) by using the following program: 95 °C for 2 min, 30 cycles of 95 °C for 30 s, 64 °C for 30 s, 72 °C for 1 min, and a final step of 72 °C for 5 min. PCR products were separated by denaturing PAGE, visualised by autoradiography using a Typhoon Trio plus scanner (GE Healthcare) and quantified using Quantity One software (Bio-Rad).

#### Western Blot Analysis

Cells were washed with PBS and lysed in lysis buffer (40 mM HEPES pH 7.4, 150 mM NaCl, 1% Triton X-100, 1 mM phenylmethylsulfonylfluoride, 1 mM sodium orthovanadate and 0.5 mM DTT) supplemented with phosphatase inhibitor and EDTA-free protease inhibitor cocktails (Roche). The concentration of total protein in cleared lysates was measured by Bradford assay and ~20 μg of each sample was analysed by western blotting followed by immunostaining using antibodies against SART1, PRPF8, Snu114, PRPF31 (against its N-terminus or C-terminus), PRPF4 and SF3B155, and the Amersham ECL detection kit (GE Healthcare). All antibody details are shown in **Table S6.**

#### Analysis of snRNP Levels by Glycerol Gradient Fractionation

Nuclear extracts (200 μg each) were diluted with an equal volume of gradient buffer (G150: 20 mM HEPES pH 7.9, 150 Mm NaCl, 1.5 mM MgCl_2_ and 0.5 mM DTT) and sedimented on linear 4 ml 10-30% (v/v) glycerol gradients in the G150 buffer. After ultracentrifugation in a Sorvall TH-660 rotor for 14 hrs at 29,000 rpm, the gradients were separated into 24 fractions. To analyse the relative levels of snRNPs in the nuclear extracts, proteins in the gradient fractions were digested by Proteinase K in 20 mM HEPES pH 7.9, 150 mMNaCl, 10 mM EDTA, 1 % (w/v) SDS for 45 min at 42 °C, the RNAs were extracted by phenol/chloroform/isoamylalcohol and precipitated. The isolated RNAs were separated by denaturing 8% urea PAGE followed by Northern blotting using 5’-end radiolabeled DNA probes against U1, U2, U4, U6 and U5 snRNAs. To analyse the association of selected splicing proteins with the tri-snRNP, proteins were precipitated from gradient fractions and separated on NuPAGE 4–12% Bis—Tris gels (Invitrogen) followed by blotting and immunostaining using antibodies against PRPF8, Brr2, Snu114, PRPF31 (against its C-terminus), PRPF4 and SF3b155, and the Amersham ECL detection kit (GE Healthcare). All antibody details are shown in **Table S6.**

### QUANTIFICATION AND STATISTICAL ANALYSIS

p-values were calculated of normally distributed data sets using a two-tailed Student’s t test, or One-way ANOVA with Dunnett’s post-hoc test, or Two-way ANOVA with Bonferroni post-hoc tests using GraphPad Prism Software Inc. (San Diego, CA, USA). Statistical analyses represent the mean of at least three independent experiments, error bars represent standard error of mean (s.e.m.) or as otherwise indicated. The statistical significance of pairwise comparisons shown on bar graphs is indicated by: n.s. not significant, * p<0.05, ** p<0.01, *** p<0.001, and **** p<0.0001. For cell populations, a minimum of 100 cells were counted from >10 separate fields of view.

### DATA AND SOFTWARE AVAILABILITY

#### Data resources

The trimmed FASTQ data for all human samples were uploaded to the European Nucleotide Archive under the accession number PRJEB22885.

***include mouse data accession number when known***

